# Cas12a2 elicits abortive infection via RNA-triggered destruction of double-stranded DNA

**DOI:** 10.1101/2022.06.13.495973

**Authors:** Oleg Dmytrenko, Gina C. Neumann, Thomson Hallmark, Dylan J. Keiser, Valerie M. Crowley, Elena Vialetto, Ioannis Mougiakos, Katharina G. Wandera, Hannah Domgaard, Johannes Weber, Josie Metcalf, Matthew B. Begemann, Benjamin N. Gray, Ryan N. Jackson, Chase L. Beisel

## Abstract

Bacterial abortive infection systems limit the spread of foreign invaders by shutting down or killing infected cells before the invaders can replicate^1, 2^. Several RNA-targeting CRISPR-Cas systems (e.g., types III and VI) cause Abi phenotypes by activating indiscriminate RNases^3–5^. However, a CRISPR-mediated abortive mechanism that relies on indiscriminate DNase activity has yet to be observed. Here we report that RNA targeting by the type V Cas12a2 nuclease drives abortive infection through non-specific cleavage of double-stranded (ds)DNA. Upon recognition of an RNA target with an activating protospacer-flanking sequence, Cas12a2 efficiently degrades single-stranded (ss)RNA, ssDNA, and dsDNA. Within cells, the dsDNase activity induces an SOS response and impairs growth, stemming the infection. Finally, we harnessed the collateral activity of Cas12a2 for direct RNA detection, demonstrating that Cas12a2 can be repurposed as an RNA-guided, RNA-targeting tool. These findings expand the known defensive capabilities of CRISPR-Cas systems and create additional opportunities for CRISPR technologies.

## MAIN TEXT

All forms of life employ defense strategies that cause cells to go dormant or die to limit the spread of infectious agents^1^. In bacteria and archaea, this strategy is called Abortive infection (Abi), and it is used by a vast variety of bacterial defense systems^1, 2^. Recently, it was shown that CRISPR RNA (crRNA)-guided adaptive immune systems that target RNA cause Abi phenotypes^3–6^. In type III systems, target RNA binding triggers production of cyclic oligoadenylate secondary messengers that in turn activate indiscriminate accessory RNases (e.g., Cms6/Csx1)^4, 5, 7–9^. Type VI systems also non-specifically degrade RNA, but instead of relying on a secondary messenger and an accessory protein, the Cas13 single-effector nuclease acts as both crRNA-guided effector and indiscriminate nuclease^3, 10, 11^. In each case, the indiscriminate RNase activities cleave both viral and host RNAs, reducing virus replication and causing cellular dormancy as an abortive phenotype^3–5^. It has been proposed that CRISPR-mediated DNase activity may also cause Abi through indiscriminate dsDNases (e.g., NucC) activated by type III system secondary messengers^12, 13^, or indiscriminate ssDNase activity from type V Cas12a effectors^14^. However, type III CRISPR-mediated dsDNase activity has yet to be examined *in vivo*, and the ssDNase activity of Cas12a has not been shown to cause Abi^15^.

Here we reveal that Cas12a2, a type V single-effector CRISPR-associated (Cas) nuclease, induces an Abi phenotype when challenged with plasmids complementary to crRNA-guides. Biochemical assays with recombinant protein revealed that Cas12a2 recognizes RNA targets that unleash non-specific dsDNA, single-stranded (ss)DNA, and ssRNA nuclease activities distinct from those of other single-subunit RNA-targeting (e.g., Cas13a) and dsDNA-targeting (e.g., Cas12a) Cas nucleases^11, 16, 17^. Additionally, we show that the Cas12a2 non-specific nuclease activities damage bacterial DNA, triggering the SOS response and impairing cell growth. Collectively these results suggest that the dsDNase activity of Cas12a2 is instrumental in triggering the Abi phenotype. Additionally, as a proof-of-principle demonstration, we show that Cas12a2 can detect RNA at a sensitivity comparable to that of RNA-targeting Cas13a nuclease at various temperatures^18^.

### Cas12a2 targeting results in an abortive infection phenotype

Cas12a2 is a type V effector nuclease previously grouped with Cas12a sequences^16^ and later described as a Cas12a variant due to similarities in the CRISPR repeat sequence and the presence of a homologous RuvC endonuclease domain^19^ (Fig. 1a). However, in our phylogenetic analysis with additional orthologs, we found that Cas12a2 sequences form a distinct clade within the type V phylogeny (Fig. 1a **and** Extended Data Fig. 1). Although Cas12a2 and Cas12a possess related RuvC domains, Cas12a2 is distinguished from Cas12a by the presence of a large domain of unknown function located in place of the Cas12a bridge helix as well as a zinc-finger domain in place of the Cas12a Nuc domain (Fig. 1b **and** Extended Data Fig. 2). Considering their original classification as well as our phylogenetic data and structural results^20^, we named these distinct type V nucleases Cas12a2.

**Fig. 1.**
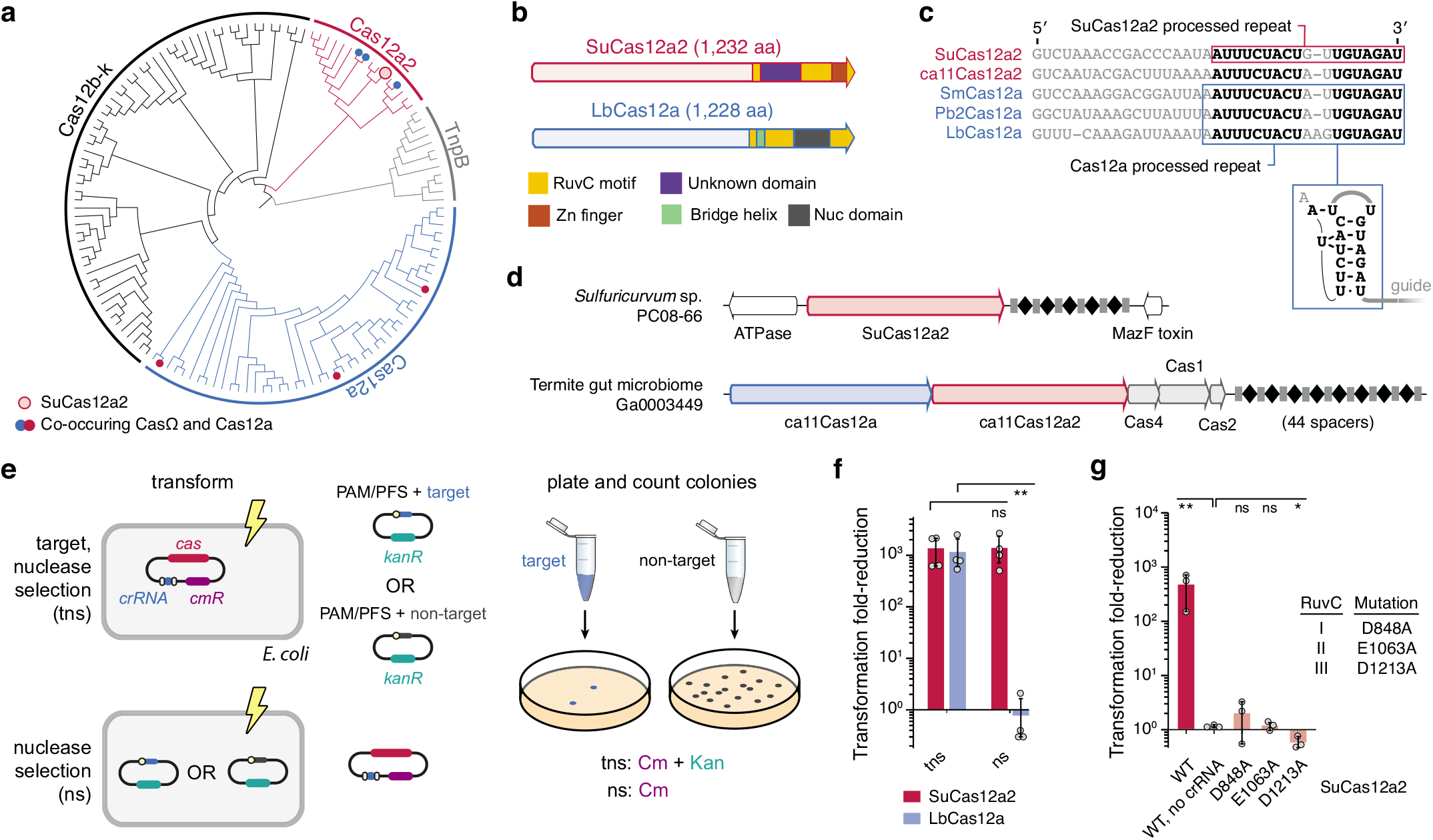
Cas12a2 nucleases form a distinct clade within type V Cas12 nucleases. (**a**) Maximum likelihood phylogenetic inference of selected Cas12 nucleases and identified Cas12a2 nucleases. See Extended Data Figure 1 for the detailed phylogeny. Systems with co-occurring Cas12a2 and Cas12a are indicated with filled red and blue circles. SuCas12a2 is indicated with an unfilled red circle. (**b**) Domain architecture of SuCas12a2 in comparison to LbCas12a. (**c**) Aligned direct repeats associated with representative Cas12a2 and Cas12a nucleases. Bolded nucleotides indicate conserved positions within the processed repeats for both nucleases. The predicted pseudoknot structure of the Cas12a repeat is diagrammed below. The loop of the hairpin (shown in gray) is variable. See Extended Data Figure 7 for pre-crRNA processing by SuCas12a2. (**d**) CRISPR system gene organizations within representative genomic loci encoding Cas12a2. Examples of systems encoding Cas12a2 as the sole Cas nuclease and also Cas12a are shown. (**e**) Diagram of the traditional (top - tns) and revised (bottom - ns) plasmid interference assay. (**f**) Reduction in plasmid transformation for SuCas12a2 and LbCa12a2 under target plasmid and nuclease plasmid selection. (**g**) Reduction in plasmid transformation of SuCas12a2 RuvC mutants under target and nuclease selection.

Intriguingly, some CRISPR-Cas systems contain both *cas12a2* and *cas12a* genes in tandem next to a shared CRISPR array (Fig. 1c). Considering that *cas12a* and *cas12a2* are adjacent to CRISPR repeats with similar sequences (Extended Data Fig. 3) and that Cas12a2 is predicted to be structurally similar throughout regions known in Cas12a to bind and process the crRNA^19^, we hypothesized that both proteins bind and process similar crRNA guides. However, because the proteins are diverse in other domains, we further hypothesized Cas12a2 performs a defense function distinct from the dsDNA-targeting activity of Cas12a^16^.

To test these hypotheses, we transferred the *cas12a2* gene from the sulfur-oxidizing epsilonproteobacterium *Sulfuricurvum* sp. PC08-66 (SuCas12a2) along with a CRISPR array on an expression plasmid into *E. coli* cells. We then performed a traditional plasmid interference assay that depletes cells by selecting for both the plasmid containing the nuclease and crRNA and a target plasmid (Fig. 1e). This assay detects broad immune system activity but cannot distinguish between defense activities that only deplete the target from those that activate Abi phenotypes. To test if Cas12a2 utilizes an Abi mechanism, we modified the plasmid interference assay to only deplete cells under Abi phenotypes by only selecting for the nuclease plasmid (Fig. 1e). Consistent with our hypothesis that Cas12a2 functions differently than Cas12a, Cas12a2 depleted cells in both the traditional (∼1,900 fold reduction) and modified (∼1,300 fold reduction) plasmid interference assays with different targeting guides, while Cas12a from *Lachnospiraceae bacterium* (LbCas12a) only depleted cells in the traditional assay (Figs 1f). Similar trends were observed with different target sites, different Cas12a2 homologs, and when assessing the Cas12a from *Prevotella bryantii B14* (Pb2Cas12a) (Extended Data Fig. 4). Additionally, mutating predicted active residues within any of the three RuvC motifs in SuCas12a2 eliminated immune function (Fig. 1g). Collectively, these results indicated that Cas12a2 relies on a RuvC nuclease domain and induces Abi with a mechanism distinct from that of Cas12a.

### RNA targeting by Cas12a2 triggers non-specific degradation of double-stranded DNA, single-stranded DNA, and single-stranded RNA

CRISPR systems that cause Abi phenotypes (e.g., types III and VI) rely on indiscriminate RNases activated by RNA targeting^3–5^. To determine if Cas12a2 utilizes a similar mechanism, we recombinantly expressed and purified SuCas12a2 and tested its enzymatic activities *in vitro* (Fig. 2 **and** Extended Data Fig. 5). However, before examining the nucleic acid targeting activities, we needed to determine how Cas12a2 crRNAs are processed.

**Fig. 2.**
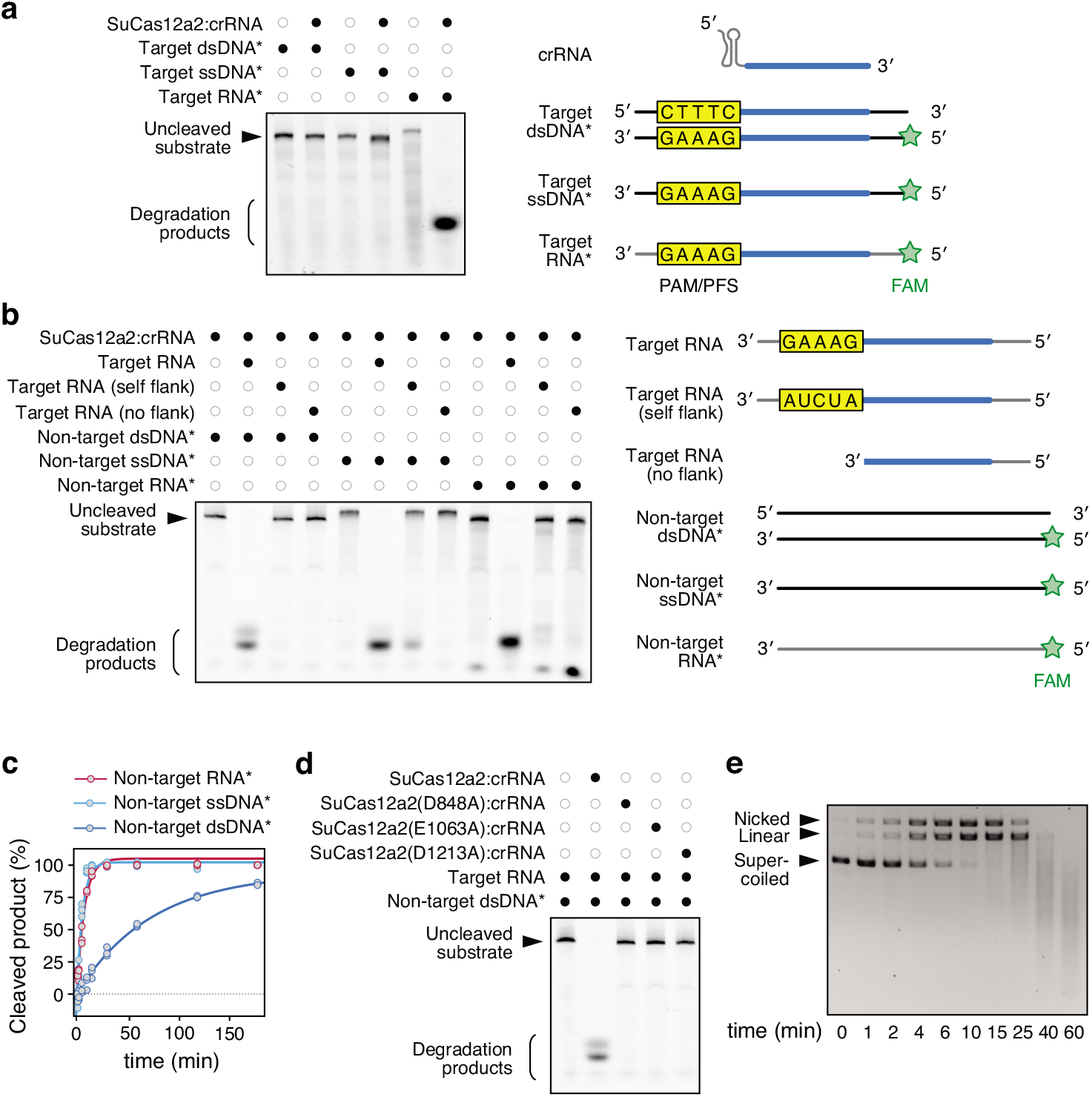
RNA target recognition by SuCas12a2 triggers degradation of ssRNA, ssDNA and dsDNA *in vitro*. (**a**) Direct targeting of different FAM-labeled nucleic-acid substrates by a purified SuCas12a2:crRNA complex, * indicates a FAM-labeled substrate, and diagrams to the right indicate substrates. (**b**) Collateral cleavage of FAM-labeled non-target nucleic-acid substrates by the SuCas12a2:crRNA complex with different target RNA substrates after 1 hour. Target RNA: non-self flanking sequence on the 3′ end. Self flank: flanking sequence mutated to the reverse complement of the crRNA repeat tag. No flank: only the reverse complement of the crRNA guide. Diagrams of target RNAs and non-target nucleic acids are diagrammed to the right. (**c**) Time course of RNA-triggered collateral cleavage of labeled non-target RNA, ssDNA, or dsDNA. See Extended Data Figure 9b for representative gel images. (**d**) Impact of mutating each of the three RuvC motifs on RNA-triggered collateral cleavage of dsDNA. All results are representative of three independent experiments. (**e**) Time course of RNA-triggered collateral cleavage of non-target plasmid DNA. Plasmid DNA was visualized with ethidium bromide.

The CRISPR repeats of Cas12a and Cas12a2 systems are highly conserved on the 3′ side (Extended Data Fig. 3), and sequence alignments predict Cas12a2 shares secondary structure in the region of the Cas12a pre-crRNA processing active site^19, 21^ (Extended Data Fig. 6). Consistent with this prediction, an RNA-processing assay coupled to RNA sequencing revealed that SuCas12a2 processes its pre-crRNAs one nucleotide (nt) downstream of the position cleaved by Cas12a. Mutating basic amino acids (K784 and R785) located in the predicted RNA processing active site abolished activity^22^ (Fig. 1c **and** Extended Data Fig. 7a-c). Similar processing was observed *in vivo* with co-expressed Cas12a2 and a single-spacer CRISPR array, producing ∼42-nt crRNAs with guide sequences ∼24 nts long (Extended Data Fig. 7d). Furthermore, plasmid interference assays revealed that Cas12a and Cas12a2 can interchange guides without impairing immunity (Extended Data Fig. 8). Therefore, the Cas12a2 nuclease processes its own crRNA guides like other type V effector nucleases^21, 23^ and can share crRNAs with Cas12a.

To determine the nucleic-acid target preference of crRNA-guided Cas12a2, complementary ssRNA, ssDNA, and dsDNA substrates containing an A/T-rich flanking sequence (paralleling Cas12a substrates^16, 22^) was fluorescently labeled with a FAM molecule and combined with crRNA-guided Cas12a2 (Fig. 2a). Similar to CRISPR-Cas systems that cause Abi, yet unlike the dsDNA-targeting Cas12a, Cas12a2 is activated only in the presence of complementary RNA targets. Additionally, this observation suggests that spurious transcription of our target plasmid is sufficient to activate the immune system *in vivo* (Fig. 1f), consistent with what has been observed with type III systems^24^.

As other Cas Abi mechanisms rely on collateral indiscriminate RNase activity, we decided to explore whether specific RNA targeting by Cas12a2 induces indiscriminate nuclease activity. We found that SuCas12a2 robustly degraded FAM-labeled ssRNA, ssDNA and dsDNA substrates; in contrast, other Cas nucleases non-specifically degrade only ssRNA (Cas13a)^11^ or ssRNA and ssDNA (Cas12g)^25^ upon RNA targeting, or only ssDNA upon dsDNA targeting (Cas12a)^14^ (Fig. 2b **and** Extended Data Fig. 9a). Of the three collateral substrates, ssRNA and ssDNA were more efficiently cleaved than ssDNA by Cas12a2 (Fig. 2c **and** Extended Data Fig. 9b). Also, similar to Cas13a, complementary ssDNA and dsDNA do not activate any Cas12a2 non-specific nuclease activity (Extended Data Fig. 9a), while dsRNA is not a substrate of collateral cleavage (Extended Data Fig. 9c).

To examine if Cas12a2 activity is reliant on detecting a ‘non-self’ signal adjacent to the target (called a protospacer flanking signal or PFS)^11^, we performed *in vitro* cleavage assays in which target RNA sequences were flanked on the 3′ side with a ‘self’ sequence complementary to the crRNA repeat (5′-AUCUA-3′), the ‘non-self’ PFS used in our *in vivo* assay (5′-GAAAG-3′), or a ‘flankless’ RNA complementary to the guide region of the crRNA, but containing no PFS (Fig. 2b). Interestingly, only the RNA target containing the ‘non-self’ PFS activated collateral nuclease activity, demonstrating that specific nucleotides on the 3′ end of the RNA target are essential for activating the collateral activity of Cas12a2. Thus, Cas12a2 uses a “non-self activation” mechanism distinct from RNA-targeting systems that rely on tag:anti-tag “self deactivation” mechanisms^11, 26, 27^. Additionally, introducing disruptive mutations to any of the three RuvC motifs or conserved cysteines within the putative Zinc finger domain abolished all non-specific cleavage (Fig. 2d **and** Extended Data Fig. 9d), consistent with our *in vivo* plasmid interference results (Fig. 1g).

Our biochemical assays demonstrated Cas12a2 could quickly remove a FAM label from linear dsDNA substrates, but it was unclear if Cas12a2 could degrade DNA lacking available 5′ or 3′ ends, such as supercoiled dsDNA. We therefore challenged crRNA-guided Cas12a2 with an RNA target and a supercoiled pUC19 plasmid. Importantly, pUC19 does not contain any sequence complementary to the Cas12a2 crRNA guide. We observed that, in the presence of target RNA, SuCas12a2 rapidly nicked, linearized, and degraded pUC19 DNA, with the DNA fully degraded within 60 minutes (Fig. 2e). This rapid destruction of supercoiled plasmid contrasts with the slow and incomplete linearization of plasmid DNA by Cas12a nucleases^28^. These data suggest a mechanism where activated SuCas12a2 is able to robustly hydrolyze the phosphodiester backbone of non-specific DNA whether it is supercoiled, nicked or linear. Comparison to Cas12a (dsDNA targeting with collateral ssDNase), Cas13a (ssRNA targeting with collateral ssRNase), and Cas13g (ssRNA targeting with collateral ssRNase and ssDNase) demonstrated that the RNA-targeting ssRNase, ssDNase, and dsDNase are unique to SuCas12a2 (Extended Data Fig. 9a). Collectively, these *in vitro* results reveal that crRNAs guide SuCas12a2 to RNA targets, activating RuvC-dependent cleavage of ssRNA, ssDNA, and dsDNA, with these activities in part or in total potentially underlying the Abi phenotype.

#### Cas12a2 exhibits PFS and targeting flexibility while evading Cas12a anti-CRISPR proteins

Although our *in vitro* data indicated an underlying mechanism for the Cas12a2 Abi phenotype, we wanted to understand in greater depth the targeting limitations of these distinct enzymes. In particular, we wanted to better understand the stringency of non-self PFS sequence recognition and penalties for mismatches between the crRNA and target. We therefore challenged SuCas12a2 with a library of plasmids encoding all possible 1,024 flanking sequences on the 3′ side of the RNA-target to the −5 position (Extended Data Fig. 10a). We found that SuCas12a2 depleted approximately half of all sequences in the library, suggesting a PFS recognition mechanism more stringent than that of Cas13, but still more promiscuous than that of most DNA-targeting systems (Fig. 3a). The depleted sequences were generally A-rich in line with a 5′-GAAAG-3′ PFS but could not be fully captured by a single consensus motif (Fig. 3a **and** Extended Data Fig. 10b-c). We further validated individual depleted sequences, including representatives within five unique motifs recognized by SuCas12a2 but not by Pb2Cas12a known among Cas12a nucleases for flexible PAM recognition (Fig. 3b **and** Extended Data Fig. 10)^29^. Consistent with its function as an RNA-targeting nuclease, the recognized sequences were broad but did not follow the expected profile if Cas12a2 is principally evaluating tag:anti-tag complementarity. These results further supported a mechanism where PFS recognition by SuCas12a2 operates similar to type III systems that require recognition of a PFS or rPAM (RNA PAM) to activate^30–33^ and distinct from the evaluation of tag:anti-tag complementarity used by RNA-targeting Cas13^11, 34^ and other type III CRISPR-Cas systems^26, 27^.

**Fig. 3.**
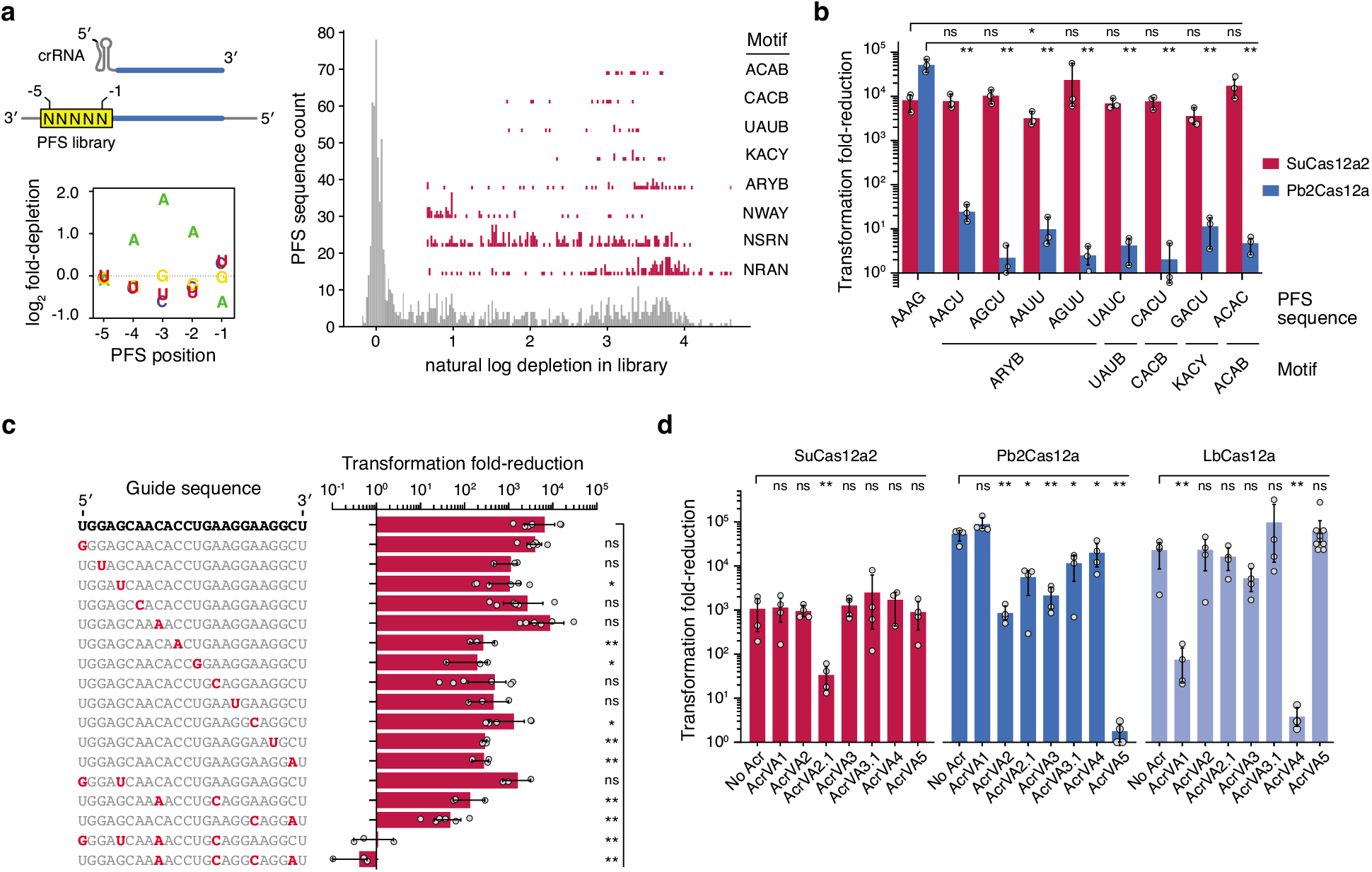
SuCas12a2 exhibits promiscuous targeting and resists anti-Cas12a proteins. (**a**) Experimentally determined PFSs and motifs recognized by SuCas12a2 in *E. coli*. Motifs capture position −4 through −1 of the PFS and are written 3′ to 5′. B = C/G/U. K = G/U. R = G/A. W = A/U. Y = C/U. Results are representative of two independent screens (see Extended Data Figure 10). (**b**) Validation of selected PFSs identified in the screen and associated with SuCas12a2 but not Pb2Cas12a. (**c**) Effect of guide mismatches on plasmid targeting by SuCas12a2 in *E. coli*. (**c**) Effect of swapping direct repeats associated with SuCas12a2 and different Cas12a nucleases. Error bars represent the mean and one standard deviation of at least three independent experiments starting from separate colonies. ns: not significant. *: p < 0.05. **: p < 0.005. (**d**) Extent of inhibition by known AcrVA proteins against SuCas12a2. Acrs were confirmed to exhibit inhibitory activity against different Cas12a homologs in *E. coli* or in cell-free transcription-translation reactions (see Extended Data Figure 12). Error bars represent the mean and one standard deviation of at least three independent experiments starting from separate colonies. ns: not significant. *: p < 0.05. **: p < 005.

Most DNA and RNA-targeting Cas nucleases have shown high sensitivity to mismatches within a seed region, where a single mismatch between the crRNA guide and target disrupts binding^11, 17, 35–37^. Therefore, to identify if SuCas12a2 relies on a seed region, we evaluated how SuCas12a2 tolerates mismatches in our cell-based assay (Fig. 3c). Interestingly, SuCas12a2 accommodated single and double mismatches across the target, with PFS-distal mutations exerting more adverse effects on plasmid targeting. To completely disrupt SuCas12a2 targeting, four mismatches were needed throughout the guide (Fig. 3c) or up to 10 mismatches on the 3′ end of a 24-nt guide (Extended Data Fig. 11). The flexible PFS recognition and a tolerance for guide:target mismatches indicates SuCas12a2 exhibits promiscuous target recognition and appears to lack a canonical seed.

Collectively, the distinct activities of Cas12a2 compared to Cas12a suggested that tandem systems possessing both nucleases may act cooperatively to broaden the effectiveness against foreign viruses and plasmids. In particular, we hypothesized that the unique structural features of Cas12a2 might prevent the escape of viruses encoding anti-CRISPR proteins that block Cas12a function^38–40^. Consistent with this hypothesis, only one (AcrVA2.1) of seven Cas12a anti-CRISPRs was able to impair Cas12a2 function, albeit only partially (Fig. 3d **and** Extended Data Fig. 12a-b). Notably, AcrVA5 also exhibited no inhibitory activity despite SuCas12a2 possessing the conserved lysine chemically modified by this Acr to block PAM recognition^41^ (Fig. 3d **and** Extended Data Fig. 12c). The limited ability of Cas12a Acrs to inhibit SuCas12a2 further underscores the distinct properties of these nucleases and the capacity of Cas12a and Cas12a2 to complement each other in nature.

#### Triggered Cas12a2 enacts widespread DNA damage and impairs growth

Although our initial results indicate Cas12a2 causes an Abi phenotype, it was unclear if the Abi mechanism causes cell dormancy or cell death. Recently it was shown that, upon recognizing an RNA target, Cas13a mediates widespread RNA degradation that drives cellular dormancy and suppresses phage infection^3^. Introducing Cas13a from *Leptotrichia shahii* (LsCas13a) into our modified plasmid interference assay, we observed that LsCas13a reduced plasmid transformation similar to Cas12a2 in the absence of antibiotic selection for the target plasmid (Extended Data Fig. 13). To verify that loss of the nuclease-and crRNA-encoding plasmids does not contribute to this result, we evaluated growth of *E. coli* in liquid culture following induction of SuCas12a2, LbCas12a, or LsCas13a with a targeting crRNA under different antibiotic selection conditions, including a no-antibiotic condition. Under these conditions, both SuCas12a2 and LsCas13a suppressed culture growth even when the target plasmids were not selected for, while LbCas12a only suppressed growth in the presence of the target plasmid antibiotic (Fig. 4a).

**Fig. 4.**
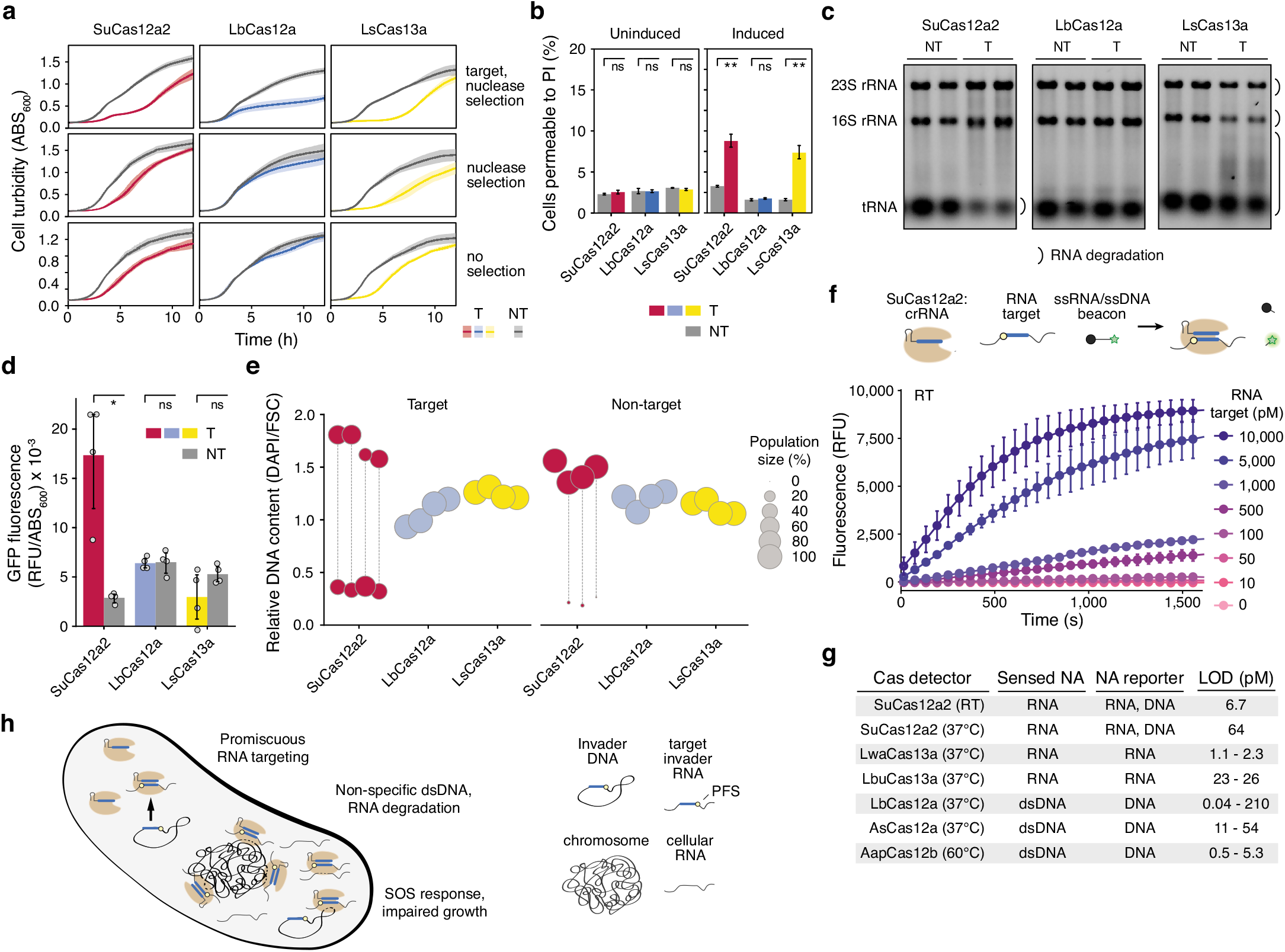
SuCas12a2 causes abortive infection principally through collateral DNA degradation and can be harnessed for RNA detection. (**a**) Reduced transformation with targeting by SuCas12a2 in the absence of target plasmid selection. The nuclease plasmid was transformed into cells with the target or non-target plasmid. (**b**) Percent of cells stained with propidium iodine indicative of loss of cell viability in the presence of a target or non-target plasmid. The Cas nuclease and associated crRNA were sampled prior to induction (uninduced) and after four hours (induced). Cells treated with isopropanol were used as a staining control. (**c**) Limited RNA degradation by SuCas12a2 compared to LsCas13a. Total RNA extracted from *E. coli* cells expressing the nuclease, crRNA, and target (T) or non-target) construct 2 hours after nuclease and crRNA induction. Duplicate independent experiments are shown. See Extended Data Figure 14 for independent quadruplicates. (**d**) Expression from an SOS-responsive GFP reporter construct in *E. coli* following four hours of plasmid targeting by SuCas12a2, LbCas12a, or LsCas13a. Targeting was conducted without antibiotic selection. (**e**) Relative DNA content in *E. coli* following four hours of plasmid targeting by SuCas12a2, LbCas12a, or LsCas13a. Targeting was conducted without antibiotic selection. DAPI fluorescence and cell size were measured by flow cytometry analysis. Each circle or pair of vertically aligned circles represents major sub-populations from the same biological replicate. (**f**) Diagram of RNA detection assay. Limit of detection for RNA detection for Cas12a2 incubated with a ssRNA beacon at room temperature (RT). Error bars in a - h represent the mean and one standard deviation of at least three independent experiments starting from separate colonies. ns: not significant. ns: not significant (p >= 0.05). *: p < 0.05. **: p < 0.005 (**g**) Unamplified limit of detection (LOD) for several Cas detectors^18^ compared to Cas12a2 determined by the velocity detection method. (**h**) Proposed model for promiscuous RNA targeting and collateral DNA degradation by SuCas12a2 and its outcome on cell growth.

Our comparison to LsCas13a suggested that the growth suppression by SuCas12a2 could occur through non-specific RNA cleavage, causing cell dormancy, while our *in vitro* data indicated that non-specific dsDNA cleavage could also suppress growth by causing cell death. To evaluate if the cells containing SuCas12a2 were undergoing cell death, we performed a cell viability assay with propidium iodide for both Cas12a2 and Cas13a. We observed only a small percentage (∼10%) of cell death in both Cas12a2 and Cas13a after 4 hours. Thus, although the indiscriminate nuclease activities of Cas12a2 cause some cell death, the primary result of Cas12a2 activity is better described as a cell dormancy phenotype (Fig. 4b).

Although Cas12a2 appears to cause dormancy, it was unclear which of the several indiscriminate nuclease activities are involved. To determine if SuCas12a2 causes dormancy through RNA cleavage, total cellular RNAs were examined under targeting and non-targeting conditions. Compared to the bacteria expressing Cas13a, Cas12a2 yielded less RNA degradation, where rRNAs appeared intact while RNAs the size of tRNAs were depleted (Fig. 4c **and** Extended Data Fig. 14).

Given the less extensive RNA degradation under targeting conditions, we asked if the indiscriminate dsDNase activity of SuCas12a2 was detectable in the context of the Abi phenotype. We reasoned that widespread dsDNA damage caused by SuCas12a2 would trigger an SOS response, impairing growth^42, 43^. In line with this assertion, plasmid targeting with SuCas12a2 but neither LbCas12a nor LsCas13a significantly induced GFP expression from an SOS-responsive reporter construct^44^ compared to a non-target control (Fig. 4d **and** Extended Data Fig. 15). Furthermore, SuCas12a2-targeting cultures diverged into two sub-populations in the absence of antibiotic selection: one represented by compact cells with reduced DNA content and another represented by filamentous cells with high DNA content (Fig. 4e **and** Extended Data Fig. 16-17). Cultures expressing LbCas12a and LsCas13a did not exhibit noticeable differences in cell sizes and DNA content for the target and non-target plasmids (Fig. 4e **and** Extended Data Fig. 16-17). Previous studies with other CRISPR-Cas systems that specifically targeted the bacterial chromosome observed similar morphological changes^45–47^, suggesting these distinct morphologies are due to dsDNA damage. These results demonstrate that RNA targeting by SuCas12a2 causes dsDNA damage of the bacterial chromosome that in turn induces the SOS response and Abi in bacteria, reflecting a distinct mechanism of immunity that relies on indiscriminate dsDNase activity. Consistent with this observation, recent cryo-EM structures reveal Cas12a2 binds and cuts dsDNA with a mechanism completely distinct from all other CRISPR associated nucleases, while structure-guided mutants with impaired *in vitro* collateral dsDNase but not ssRNase and ssDNase activities abolished *in vivo* defense activity^20^.

#### Cas12a2 can be repurposed for programmable RNA detection

CRISPR single-effector nucleases have been repurposed for a myriad of applications from gene editing to molecular diagnostics. To determine if Cas12a2 could be repurposed as a biotechnological tool, we programmed SuCas12a2 to detect RNA. We programmed apo-SuCas12a2 with a crRNA guide complementary to an RNA target and incubated the complex with a ssDNA or ssRNA beacon that fluoresces upon cleavage due to separation of a fluorophore and a quencher (Fig. 4f). Using this approach, we were able to detect RNA using both ssDNA and RNA probes at 37°C and room temperature, with a limit of detection within the range observed for other single-subunit Cas nucleases^18^ (Fig. 4f **and** Extended Data Fig. 18). These data indicate that SuCas12a2 can be readily repurposed as a tool for applications in science and medicine. We anticipate that the unique activities of this enzyme can be further leveraged to expand the CRISPR-based toolkit.

## Discussion

Collectively, our data support a model in which Cas12a2 nucleases exhibit RNA-triggered degradation of cytoplasmic dsDNA and to some extent RNA, impeding host cell growth and eliciting an Abi phenotype (Fig. 4h). This mechanism contrasts with targeted invader clearance or Abi activities exhibited by other CRISPR-Cas systems. Specifically, the mechanism exhibited by Cas12a2 is reminiscent of the recently described CBASS defense system, which relies on the indiscriminate double-stranded DNAse NucC that degrades the host cell DNA and kills the cell^12^. Interestingly, some type III systems encode NucC enzymes, suggesting that other CRISPR-Cas systems have convergently evolved to use a similar DNA degrading Abi mechanism^13, 48^.

In addition to damaging the genome and inducing the SOS response, SuCas12a2 exhibits promiscuous RNA recognition through a flexible PFS and mismatch tolerance. This flexibility could be particularly advantageous against rapidly evolving phages and could complement precise recognition and clearance of DNA targets by Cas12a in organisms that encode both Cas12a and Cas12a2 adjacent to a single CRISPR array (Fig. 1 **and** Extended Data Figure 1). This dual-nuclease strategy would be akin to bacteria encoding multiple CRISPR-Cas systems targeting the same invader^49^. However, more work is needed to understand how these two nucleases work together to counter infections.

The combination of nuclease-mediated crRNA biogenesis, RNA targeting, and collateral cleavage of ssRNA, ssDNA, and in particular dsDNA sets Cas12a2 apart from other known Cas nucleases. The apparent need to recognize the A-rich flanking sequence by SuCas12a2 to activate the indiscriminate RuvC nuclease activity strongly indicates that Cas12a2 must bind a correct PFS adjacent to the RNA target to activate cleavage rather than rely on complementarity between the repeat tag and target anti-tag pair to distinguish self from non-self sequences typical of several other RNA-targeting Cas nucleases and complexes^11, 26, 27, 50^. Investigating the underlying molecular basis of target recognition and activation of collateral cleavage by Cas12a2 could reveal new mechanisms employed by CRISPR nucleases for discriminating self from non-self targets. Recent cryo-EM structures of Cas12a2 at stages of RNA-targeting and collateral dsDNA capture are already fulfilling this need^20^.

Cas12a2 holds substantial potential for CRISPR technologies. As a proof-of-principle demonstration, we showed that SuCas12a2 can be repurposed for RNA detection with a limit of detection comparable to existing single-effector based tools^18^. Beyond the ability to detect RNA, we envision a variety of SuCas12a2 applications that expand and enhance the CRISPR-based tool kit. RNA-triggered dsDNA cleavage could allow for programmable killing of prokaryotic and eukaryotic cells with various applications, including programmable shaping of microbial communities, cancer therapeutics, and counterselection to enhance genome editing. Additionally, the ability of Cas12a2 and Cas12a to utilize the same crRNA sequence yet recognize distinct nucleic acid species (RNA versus ssDNA and dsDNA) and elicit distinct non-specific cleavage activities (ssRNA, ssDNA, and dsDNA (Cas12a2) versus ssDNA (Cas12a)) could augment existing Cas12a applications by incorporating Cas12a2. By further exploring the properties of SuCas12a2 and its homologs, we expect the advent of new and improved CRISPR technologies that could broadly benefit society.

## ACKNOWLEDGMENTS

We thank Kira Makarova for guidance naming Cas12a2, Leonhard Fläxl for assistance with TXTL, Fani Ttofali for assistance with plasmid cloning and testing, April Pawluk and Ahsen Özcan for critical feedback on the manuscript. Plasmids pET28a-mH6-Cas12g1 (Addgene plasmid # 120879) and pACYC-Cas12g1 (Addgene plasmid # 120880) were gifts from Arbor Biosciences. Plasmid pC001 was a gift from Feng Zhang (Addgene plasmid # 79150).

## Author contributions

Conceptualization: O.D., G.C.N., R.N.J., and C.L.B.; Methodology: O.D., G.D. V.M.C., J.M., H.D., T.H., and D.K. Investigation: bioinformatic discovery of Cas12a2 orthologs - B.N.G. and O.D.; first observation of bacterial Abi: G.C.N.; phylogenetic analysis - O.D.; discovery of RNA targeting: O.D., D.K., R.N.J., and C.L.B.; experimentation in *E. coli* - O.D., G.C.N., V.M.C., E.V., I.M., and J.W.; in TXTL - K.W. and J.W.; *in vitro* - V.M.C., D.K., H.D, T.H. and J.M.; Writing – Original Draft: O.D., V.M.C., R.N.J., and C.L.B.; Writing – Review and Editing: All authors; Visualization: O.D., D.K., T.H., H.D., R.N.J., and C.L.B.; Supervision: M.B.B., R.N.J., and C.L.B.; Funding Acquisition: R.N.J. and C.L.B.

## Data and materials availability

The NGS data from the PAM depletion assay and crRNA sequencing data were deposited to NCBI GEO under the accession GSE178536. All other data in the main text or the supplementary materials are available upon reasonable request.

To review GEO accession GSE178536 prior to publication, go to https://www.ncbi.nlm.nih.gov/geo/query/acc.cgi?acc=GSE178536 and enter the token mhuzyayihxyfncf.

## EXTENDED DATA

### MATERIALS AND METHODS

#### Identification of the putative Cas12a2 nucleases

Several Cas12a2 sequences were initially identified and tentatively classified as encoding Cas12a nucleases^16^. These Cas12a2 protein sequences were used as seeds for BLASTP searches of protein data in NCBI and for TBLASTN searches of metagenomic data in NCBI (https://www.ncbi.nlm.nih.gov) and JGI (https://img.jgi.doe.gov) to identify additional putative Cas12a2 nucleases.

#### Phylogenetic analysis of Cas12a2 proteins within type V systems

Amino acid sequences of Cas12a2 orthologs, Cas12a nucleases, representative nucleases from all known type V subtypes, and TnpB orthologs were aligned using Clustal Omega^51^. The resulting alignment was used to create a maximum likelihood phylogeny using RAxML-NG^52^ with the following parameters: --model JTT+G --bs-metric fbp, tbe --tree pars{60}, rand{60} --seed 12345 --bs-trees autoMRE. TnpB sequences were used as an outgroup. The amino acid sequences used in the creation of the phylogeny can be found in Extended Data File 1.

#### Domain annotation and structure prediction

Conserved motifs in SuCas12a2 were identified using MOTIF Search (https://www.genome.jp/tools/motif/, accessed on 2021.06.15) and Phyre 2^53^ (accessed on 2021.03.08). HHpred secondary structure predictions of Cas12a2 orthologous amino acid sequences were performed to identify common secondary structure between Cas12a2 and Cas12a that predicted the crRNA processing site of Cas12a2^54^.

#### Strains and plasmids

All of the *in vivo* experiments, unless indicated otherwise, were performed in *E. coli* BL21(AI). For propagation the cultures were grown in LB medium at 37°C with constant shaking at 225 - 250 rpm. *E. coli* strain TOP10 was used for plasmid cloning (**Extended Data Table 1, Tab 1**). All primers, gBlocks, and oligos were obtained from Integrated DNA Technologies, unless specified otherwise. Gibson assembly of plasmid construction was performed using NEBuilder HiFi DNA Assembly Master Mix (New England Biolabs, E2621). Mutagenesis of the plasmids, including small insertions and nucleotide substitutions, were done with Q5 Site-Directed Mutagenesis Kit (New England Biolabs, E0554S). All of the nucleases together with crRNA, unless specified otherwise, were expressed from plasmids containing p15A origin-of-replication and a chloramphenicol resistance marker. The expression of the nucleases and crRNA was controlled by a T7 promoter, unless specified otherwise. All target and non-target plasmids were created by introducing protospacer sequences and corresponding flanking sequences into pBR322 or sc101 origin-of-replication plasmids bearing a kanamycin resistance cassette, unless specified otherwise. Sequences encoding Cas12a2 orthologs (**Extended Data File 1**) were codon optimized and synthesized by Genscript. Sequences encoding Pb2Cas12a from *Prevotella bryantii B14* (NCBI Accession: WP_039871282) LbCas12a from *Lachnospiraceae bacterium* ND2006 (NCBI Accession: WP_035635841.1), FnCas12a from *Francisella tularensis* (NCBI Accession: WP_104928540.1), AsCas12a from *Acidaminococcus sp*. BV3L6 (NCBI Accession: WP_021736722.1), and Mb3Cas13a from *Moraxella bovoculi* (NCBI Accession: WP_080946945.1)^16^ were codon optimized for expression in *E. coli* and ordered as gBlocks from Integrated DNA Technologies. Sequences encoding anti-CRISPR proteins^38, 40^ (**Extended Data Table 1, Tab 3**) were codon-optimized for expression in *E. coli* and ordered as gBlocks from Integrated DNA Technologies. The Acr genes were then PCR amplified and introduced into pBAD24 plasmid backbone carrying ampicillin resistance cassette^55^. The LsCas13a-encoding plasmid pCBS2091 was ordered from Addgene (79150)^11^. For detecting RecA-dependent SOS response in *E. coli* BL21(AI), reporter plasmids pCBS2000, pCBS3611, and pCBS3616 were created by introducing *recA* promoter, included 100 bp upstream of the predicted LexA binding site, upstream of the GFP-encoding gene into plasmid pCBS198. Plasmids pCBS3611 and pCBS3616 received an ampicillin resistance cassette from plasmid pCB672^29^. The *recA* promoter sequence was identified in the genome of *E. coli* BL21(AI) between positions 2,635,525 and 2,635,347 (NCBI Accession: CP047231.1). Control plasmids pCBS3616 and pCBS2002 without the GFP-reporter genes were generated by PCR amplification of pCBS2000 and pCBS3616 followed by KLD assembly (New England Biolabs, M0554). The full list of plasmids used in the study, including links to plasmid maps, can be found in **Extended Data Table 1, Tab 2**. Relevant oligonucleotide, dsDNA, and RNA sequences are listed in **Extended Data Table 1, Tab 3**.

### *In vitro* characterization of SuCas12a2

#### Expression and purification of SuCas12a2

N-terminal 6x His-tagged SuCas12a2 WT and mutant constructs were expressed in *E. coli* Nico21(DE3) cells from a pACYC plasmid either lacking (apo) (plasmid 1416) or containing a three-spacer CRISPR array (crRNA-guided) (plasmid 1408) using either an auto-or IPTG induction. Autoinduction growths followed guidelines in Studier^56^. Briefly, a solution containing recommended concentrations of ZY media, MgSO_4_, Metals Mix, 5052 and NPS autoinduction buffers along with antibiotics needed for selection, was inoculated with bacteria from a glycerol stock or a fresh transformation. The cells were grown for five hours at 37°C shaking at ∼250 rpm and then moved to 24°C where they were incubated for 24 h before harvesting via centrifugation at 8K RPM for 25 min. Cell pellets were then stored at −80°C until purification. For the IPTG induction, 1 L of TB media was inoculated with 20 ml of overnight growth and was grown at 37°C until an OD_600_ of 0.6. The cells were then cold-shocked on ice for 15 minutes and induced with 0.1 mM IPTG, followed by a 16-18 h incubation at 18°C. Cells were harvested by centrifugation. Cells were lysed by sonication in Lysis buffer (25 mM Tris-pH 7.2, 500 mM NaCl, 10 mM imidazole, 2 mM MgCl-_2_, 10% glycerol) in the presence of leupeptin, aprotinin, pepstatin, aebsf, and lysozyme. The lysate was clarified by centrifugation at 36,400 x g for 35 minutes. Clarified lysate was added to 5 ml of Ni-NTA resin and batch bound at 4°C for 30 minutes, and then washed with 100 ml of lysis buffer. The protein was eluted with 50 ml of Ni-elution buffer (25mM Tris-pH 7.2, 500 mM NaCl, 250 mM imidazole, 2 MgCl-_2_, 10% glycerol). Fractions containing SuCas12a2 were desalted with a Hiprep 26/10 desalting column into low-salt buffer (25mM Tris-pCas12a22, 50 mM NaCl, 2 mM MgCl-_2_, 10% glycerol). SuCas12a2 + crRNA was then applied to a Hitrap Q HP column anion exchange column while the apo SuCas12a2 was applied to a Hitrap SP HP cation exchange column. The column was washed with 10% high-salt buffer (25mM Tris-pH 7.2, 1 M NaCl, 2 mM MgCl-_2_, 10% glycerol) followed by a gradient elution to 100 percent high salt buffer 10 CV (50 ml). The fractions containing SuCas12a2 were concentrated using a 100 MWKO concentrator to about 1 ml and then purified by size exclusion column chromatography over a Hiload 26/600 superdex 200 pg column equilibrated in SEC buffer (100 mM HEPES-pH 7.2, 150 mM KCl, 2mM MgCl_2_, 10% glycerol). The fractions containing SuCas12a2 were concentrated and stored at −80°C.

#### Pre-crRNA processing

##### Processing of a 3X pre-crRNA

SuCas12a2 pre-CRISPRx3 RNA was in vitro transcribed using the HiScribe™ T7 High Yield RNA Synthesis Kit (New England Biolabs). The template DNA was derived from Jackson lab plasmid 1409 linearizing with the KpnI restriction enzyme. A contaminating band that runs approximately at 130 nts was observed to be an artefact of the reaction. Numerous strategies were attempted to prevent the transcription of this contaminating band, to no success. In vitro transcribed RNA was cleaned on RNeasy spin columns (Qiagen). 1.5 μM of apo-SuCas12a2 was incubated with 1 mg of SuCas12a2 pre-CRISPRx3 RNA in 1X3.1 Buffer from New England Biolabs (100 mM NaCl, 50 mM Tris-HCl, 10 mM MgCl_2_, 100 mg/ml BSA pH 7.9) and incubated at 25°C for various times. Samples were run on a gel (12% polyacrylamide, 8M, TBE) alongside a ssRNA low range ladder (New England Biolabs) and stained with SYBR gold (ThermoFisher Scientific).

##### Processing a 1X crRNA with WT and crRNA processing mutants

A synthetic crRNA with a 13 base 5′ unprocessed overhang (**table S1 Tab 3**, smcrRNA) was refolded using the protocol outlined in Lapinaite et al., 2020^57^. In a 10 μl reaction, 150 nM of crRNA substrate was combined with 1.5 μM WT, K784A, or K785A apo SuCas12a2 protein in NEB 3.1. The reactions were incubated at 37°C for 1 h. The reactions were quenched with phenol and phenol:chloroform extracted. The results were analyzed using 12% Urea-PAGE stained with SYBR-GOLD.

#### Nucleic acid cleavage assays

##### Analysis of targeted cleavage

10 μl reactions of 250 nM SuCas12a2:crRNA with 100 nM of complementary FAM-labeled synthetic oligonucleotide (i.e., ssDNA, dsDNA or RNA) in 1X NEB 3.1 buffer were incubated at 37°C for 1 h. Reactions were quenched with phenol and then phenol:chloroform extracted. Results were analyzed using the FDF-PAGE method outlined in Harris *et al.*,^58^ and visualized for fluorescein fluorescence.

##### Analysis of collateral cleavage

10 μl reactions of 250 nM SuCas12a2:crRNA, and 250 nM of target (RNA complementary to the crRNA-guide) or non-target (RNA non-complementary to the crRNA-guide) substrate and 100 nM of 5′-FAM labeled collateral substrate (ssDNA, dsDNA, RNA) in 1X NEB 3.1 were incubated at 37°C for 1 h. Reactions were quenched with phenol and then phenol:chloroform extracted. The results were analyzed using 12% Urea-PAGE and visualized for fluorescein fluorescence.

##### Analysis of flanking sequence requirements for activation

10 μl reactions of 250 nM Cas12a2:crRNA, with 300 nM of different target ssRNAs: Self (flanked by sequence complementary to the direct repeat of the crRNA), no flanks, and flanks containing a 5′-GAAA-3′ protospacer flanking sequence on the 3’ side of the protospacer, and 100 nM of collateral 5′-FAM dsDNA in 1X NEB 3.1 buffer were incubated at 37°C for 1 h. The reactions were quenched with phenol and phenol:chloroform extracted. The results were analyzed using 12% Urea-PAGE and visualized for fluorescein fluorescence.

##### Kinetic analysis of collateral cleavage

A single 100 μl reaction containing 100 nM Cas12a2:crRNA, 100 nM of target ssRNA (crRNA complementary) and 100 nM of different 5′-FAM labeled collateral substrates (ssDNA, dsDNA, RNA) in 1X NEB 3.1 buffer was made. Time points were taken at 1, 2, 5, 10, 15, 30, 60, 120, and 180 min by combining 10 μl from the 100 μl reaction with phenol, followed by phenol:chloroform extraction. The results were analyzed using 12% Urea-PAGE and visualized for fluorescein fluorescence.

##### Plasmid cleavage assay

A 100 μl reaction containing 14 nM Cas12a2:crRNA, 25 nM target RNA, 7 nM of pUC19 plasmid in 1X NEB 3.1 buffer was incubated at 37°C. At indicated time points, 10 μl of the reaction was removed and quenched with phenol followed by phenol:chloroform extraction. The reactions were visualized on 1% agarose with ethidium bromide.

#### Collateral cleavage comparison of Cas12a2 with Cas12a, Cas13a, and Cas12g

EnGen LbaCas12a (LbCas12a) was purchased from New England Biolabs (M0653S). 10-μl reactions containing 250 nM of LbCas12a and 500 nM of its cognate crRNA in 1X NEB 2.1 buffer were incubated at 37°C with 200 nM of different target substrates (ssDNA, dsDNA, RNA) and 100 nM of different FAM labeled collateral substrates (ssDNA, dsDNA, RNA). After 1 h, the reactions were quenched by phenol and phenol:chloroform extracted. The results were analyzed using 12% Urea-PAGE and visualized for fluorescein fluorescence.

LwCas13a was purchased from MCLAB Molecular Cloning Laboratories (Cas13a-100). 10-μl reactions containing 250 nM of LwCas13a and 500 nM of its cognate crRNA in the provided 1X Cas9 buffer (20 mM HEPES (pH 6.5), 5 mM MgCl_2_, 100mM NaCl, 100 μM EDTA) were incubated at 37°C with 200 nM of different target substrates (ssDNA, dsDNA, RNA) and 100 nM of different FAM labeled collateral substrates (ssDNA, dsDNA, RNA). After 1 h, the reactions were quenched by phenol and phenol:chloroform extracted. The results were analyzed using 12% Urea-PAGE and visualized for fluorescein fluorescence.

AbCas12g was expressed in *E. coli* NiCo 21 DE3 using pET28a-mH6-Cas12g1 (Addgene plasmid #120879) and initially purified as described previously^25^. The protein was then transferred to low salt buffer (25 mM HEPES pH 7.8, 50 mM NH_4_Cl, 2 mM MgCl_2_, 7 mM BME, 5% glycerol) by buffer exchange and loaded over heparin followed by elution with a linear NaCl gradient and gel filtration as described previously^59^. Purified protein was flash frozen and stored at −80°C. The Cas12g1 non-coding plasmid pACYC-Cas12g1 (Addgene plasmid #120880) was used as a template for PCR amplification of the AbCas12g tracrRNA sequence with Cas12gtracrRNA F and R primers (Table S1 tab 3) in 2x Taq Master Mix (New England Biolabs). The non-coding plasmid was removed with DpnI by incubation at 37°C for 1 h in CutSmart buffer (New England Biolabs). DNA components were cleaned after PCR and DpnI digest with E.Z.N.A. Cycle Pure Kit (OMEGA BioTek). The Cas12g tracrRNA was transcribed with HighScribe T7 Quick High Yield RNA synthesis kit and cleaned with Monarch RNA cleanup kit (New England Biolabs). 10-μl reactions containing 250 nM of Cas12g, 500 nM of the Cas12g crRNA, and 1 μΜ of Cas12g tracrRNA in 1X NEB 3.1 buffer were incubated at 37°C or 50°C with 200 nM of different target substrates (ssDNA, dsDNA, RNA) and 100 nM of different FAM labeled collateral substrates (ssDNA, dsDNA, RNA). After 1 h, the reactions were quenched by phenol and phenol:chloroform extracted. The results were analyzed using 12% Urea-PAGE and visualized for fluorescein fluorescence.

To analyze SuCas12a2 collateral activity, 10-μl reactions containing 250 nM of Cas12a2:crRNA, 200 nM of different target substrates (ssDNA, dsDNA, ssRNA) and 100 nM of different FAM labeled collateral substrates (ssDNA, dsDNA, RNA) in 1X NEB 3.1 buffer were incubated at 37°C for 1 h. The reactions were quenched with phenol and phenol:chloroform extracted. The results were analyzed using 12% Urea-PAGE and visualized for fluorescein.

#### RNA detection by Cas12a2 with ssRNA and ssDNA reporter probes

Cas12a2 (100nM) was complexed with crRNA (120nM) in NEB 3.1 buffer (50 mM Tris-HCl pH 7.9, 100 mM NaCl, 10 mM MgCl2, 100 µg/ml BSA) before combining with RNAse or DNAse Alert (200nM, IDT) and Target RNA to the indicated concentrations in a 384 well plate (Greiner Bio-One, Ref#. 784077). A background control was prepared with nuclease free water instead of Target RNA. Reactions were monitored for Reporter fluorescence (RNAse Alert: Ex. 485-20/ Em. 528-20, DNAse Alert: Ex. 500-20/Em. 560-20) over time at either ambient conditions (RT) or 37°C using a Synergy H4 Hybrid multi-mode microplate reader (BioTek instruments inc.). The slope of the linear region (between 5 and 30 min) was determined at each concentration of target RNA using GraphPad PRISM. Standard error of the linear fit was used as a proxy for standard deviation, and the limit of detection was calculated as 3 x Standard Error of the water background as in (ref. ^18^). Limit of detection was estimated by determining where the plot of V_O_ vs. [Target RNA] crosses the detection threshold.

### Cas12a2 characterization in *E. coli*

#### crRNA sequencing and analysis

The SuCas12a2 expression plasmid pCBS3568 containing the nuclease- and the crRNA-encoding sequences and the no-crRNA control pCBS3569 were transformed into *E. coli* BL21(AI) and the transformants were plated on selection pates. The resulting colonies were picked and used to inoculate 2 ml overnight liquid cultures. The next day, the overnight cultures were used to inoculate 25 ml of LB containing chloramphenicol to an OD_600_ of approximately 0.05. Once the growing cultures reached the OD_600_ of 0.25 after approximately 40 min, expression of the nuclease and the crRNA were induced with 1 mM isopropyl β-d-1-thiogalactopyranoside (IPTG) and 0.2% L-arabinose. The induced cultures were harvested in the stationary phase by centrifugation at 14,000 rpm and 4°C for 2 min. Afterwards, the cell pellets were immediately frozen in liquid N_2_ and stored at −80°C until further processing.

Total RNA was purified from cell pellets using Direct-zol RNA Miniprep Plus (Zymo Research, R2072) following manufacturer’s instructions. DNA was removed using Turbo DNase (Life Technologies, AM2238). Between the individual processing steps RNA was purified using RNA Clean & Concentrator kit (Zymo Research, R1017). Ribosomal RNA was removed from the samples using RiboMinus Transcriptome Isolation Kit, bacteria (ThermoFisher Scientific, K155004). 3′-phosphoryl groups were removed from RNA using T4 polynucleotide kinase (New England Biolabs, M0201S). cDNA synthesis and library preparation was performed with NEBNext Multiplex Small RNA Library Prep Set for Illumina (New England Biolabs, E7330S). Size selection for fragments between 200 bp and 700 bp was performed with Select-a-Size DNA Clean & Concentrator kit (Zymo Research, D4080). Finally, DNA was purified using AMPure XP beads (Beckman Coulter, A63882) and quantified with the Qubit dsDNA HS assay kit (ThermoFisher Scientific, Q32851) on DeNovix DS-11 FX (DeNovix).

Library sequencing was performed at the Helmholtz Center for Infectious Research (HZI) GMAK facility in Braunschweig, Germany, using the MiSeq 300 sequencing method (Illumina). The resulting paired-end reads were quality controlled, trimmed, and merged using BBTools^60^ (sourceforge.net/projects/bbmap/). Afterwards, the reads were mapped to the crRNA expression site on the plus strand of pCBS273 using Bowtie2 (http://bowtie-bio.sourceforge.net/bowtie2/). The associated raw and process sequencing data as well the data processing steps can be found on NCBI GEO (Accession: GSE178531).

#### Plasmid clearance assay in *E. coli*

Standard plasmid clearance assay was performed in *E. coli* BL21(AI) containing nuclease- and crRNA-expressing plasmids. Bacterial cultures were grown overnight and used to inoculate fresh LB medium containing chloramphenicol to an OD_600_ of 0.05 - 0.1. Subsequently, these cultures were grown until the OD_600_ reached approximately 0.25, at which time 1 mM IPTG and 0.2% L-arabinose were added for induction. Once the cultures reached the OD_600_ between 0.6 and 0.8, the cells were harvested and made electrocompetent^61^. Electrocompetent cells were prepared from four biological replicates. Immediately after, 1 µl of 50 ng/ul of the target and non-target plasmid were electroporated into 50 µl of the electrocompetent *E. coli* cells. To achieve high transformation efficiencies, the used plasmids were purified through ethanol precipitation and quantified using the Qubit dsDNA HS Assay Kit (ThermoFisher Scientific, Q32851). The electroporated cells were recovered for 1 hour at 37°C with shaking in 500 µl LB containing 1 mM IPTG and 0.2% L-arabinose without antibiotics. Afterwards, the cultures were sequentially diluted to 10^-5^ in 10-fold increments. 5-10 µl of each dilution were spotted on LB plates containing antibiotics to select the nuclease-crRNA and the target/non-target plasmids. Additionally the plates contained 0.3 mM IPTG and 0.2% L-arabinose. The plates were incubated overnight at 37°C.

The next day the colonies were manually counted and the resulting counts adjusted for the dilution factor. Counts from the highest countable dilution were used to calculate transformation fold reduction as a ratio between the colonies in the non-target condition divided by the colonies in the target condition.

In a modification of the assay used to determine the cell suicide phenotype, the target and the non-target plasmids were transformed into *E. coli* BL21(AI) first. Next, these cells were made electrocompetent and the nuclease-crRNA plasmids were transformed in last.

When testing Acrs, the Acr plasmid (ampicillin) and the nuclease-crRNA plasmid (chloramphenicol) were co-transformed, followed by electroporation of the target or non-target plasmid (kanamycin).

#### Growth experiments

To investigate growth of the cultures under nuclease targeting conditions, the nuclease-crRNA and the target/non-target plasmids were transformed into *E. coli* BL21(AI). The resulting transformants were recovered in SOC medium and grown overnight with 0.2% glucose to inhibit nuclease and crRNA expression. In the morning, the cells were harvested by centrifugation at 5,000 g for 2 min. The pellets were resuspended in LB and used to inoculate 200 µl of LB medium on a 96-well plate to the final OD_600_ of 0.01. Depending on the experiment, the reactions contained different combinations of antibiotics, IPTG, and L-arabinose. The plates were incubated in a BioTek Synergy H1 plate reader at 37°C with vigorous shaking. The OD_600_ of the cultures were recorded every 3 min. Plasmid clearance assay was performed with the overnight cultures, as described above.

#### PFS depletion assay in *E. coli*

To determine PFS preferences of SuCas12a2, a PFS depletion assay was performed. An oligo library (ODpr23) consisting of 1,024 nucleotide combinations in place of a 5 nt PFS-encoding site was synthesized by Integrated DNA Technologies. Using the ODpr23 oligo pool library in a combination with primer ODpr24, targeting plasmid pCBS276 was PCR amplified using Q5 polymerase (New England Biolabs, M0543). The PCR products were gel-purified using Zymoclean Gel DNA Recovery Kit (Zymo Research, D4007) and ligated using KLD reaction mix (New England Biolabs, M0554). The ligated plasmids were purified using ethanol precipitation and electroporated into *E. coli* TOP10. A total of 10 electroporation reactions were performed. Following recovery of the electroporated cells in SOC medium, the individual reactions were combined to inoculate 90 ml of LB medium containing kanamycin. 10 µl from each electroporation reaction were plated on selective LB medium to estimate the total number of transformed bacteria. With the colony counts we estimated that the total number of transformed cells exceeded the number of unique PAM sequences in the library (1,024) by approximately 2,300-fold. Plasmid library DNA was purified from the combined overnight culture using ZymoPURE II Plasmid Midiprep Kit (Zymo Research, D4201) and additionally cleaned by ethanol precipitation. Next, the plasmid library was verified by Sanger sequencing.

The PAM plasmid library was transformed into electrocompetent *E. coli* BL21(AI) containing either the SuCas12a2 nuclease-expressing plasmid pCBS273 and an empty plasmid control pCBS3569. The electrocompetent cells were prepared as described above. Approximately 600 ng of the plasmid DNA were electroporated into 50 µl volume of the competent cells. The transformed bacteria were recovered in 500 µl of SOC medium for 1 h at 37°C and were used to inoculate 50 ml LB with 1 mM IPTG and 0.2% L-arabinose in the presence of kanamycin and chloramphenicol. The cultures were grown for 13 hours before the cells were harvested by centrifugation at 4,000 g for 15 min and the plasmid DNA extracted using ZymoPURE II Plasmid Midiprep Kit (Zymo Research, D4201). Following recovery, bacteria were also plated on LB plates containing kanamycin and chloramphenicol without the inducers. These plates were used to estimate the total number of cells transformed with the plasmid library. The total number of transformed cells estimated based on the colony counts exceeded the number of unique PAM sequences in the library by approximately 1,700-fold for the cells containing the SuCas12a2-crRNA plasmid (pCBS273) compared to 11,900-fold in the no-crRNA control (pCBS3569).

The region of the plasmid DNA which contained the target site including the PFS-encoding sequence was PCR amplified using primers ODpr55 and ODpr56. The PCR reactions were purified using AMPure XP beads (Beckman Coulter, A63882). The purified PCR products were indexed using primers ODpr58, ODpr60, ODpr59, and ODpr61. The indexed PCR products were purified using the AMPure XP beads, quantified with the Qubit assay (ThermoFisher Scientific, Q32851), and sent for sequencing at the HZI GMAK facility in Braunschweig, Germany, using MiSeq PE300 Illumina sequencing method.

Analysis of the PFS-encoding sequences depletion data as well as the creation of the PFS wheels were performed as described previously^62^. PFS consensus motifs were defined manually. The raw and the processed sequencing data as well as the data processing steps can be found on NCBI GEO (Accession: GSE178530). Individual PFS sequences were validated using plasmid clearance assay as described above.

#### Cell-free transcription-translation (TXTL) reactions

*In vitro assay to test Acr sensitivity of Cas12a nucleases -* Plasmids encoding Cas12a nuclease were pre-expressed together with a plasmid encoding either a target or non-target crRNA in 9 µl of MyTXTL master mix (Arbor Biosciences) at the final concentration of 4 nM for each plasmid in the total volume of 12 µl. Acrs were pre-expressed separately, at the concentration of 4 nM in the total volume of 12 µl. Since the Acrs are encoded on linear DNA fragments, GamS at a final concentration of 2 µM, was added to prevent DNA degradation. All pre-expressions were carried out at 29°C for 16 hours. The subsequent cleavage assay was performed by adding 1 µl of each pre-expression reaction to 9 µl of fresh myTXTL mix. pCBS420 plasmid constitutively expressing deGFP protein was used as a reporter at the final concentration of 1 nM. For quantification, four 3 µl replicates per reaction were transferred onto a 96-well V-bottom plate (Corning Costar 3357). The reactions were prepared using Echo 525 Liquid Handler (Beckman Coulter). Fluorescence was measured on a BioTek Synergy H1 plate reader (Excitation: 485/20, Emission: 528/20). Time-course measurements were run for 16 hours at 29°C, with 3 minute intervals between the measurements.

All fold-repression values for plasmid reporter constructs represent the ratio of deGFP concentrations after 16 h of reaction for the non-target over the target crRNA. For the experiments measuring the inhibitory activity of Acrs, inhibition was calculated from endpoint expression values after 16 h of expression according to the following formula^63^.

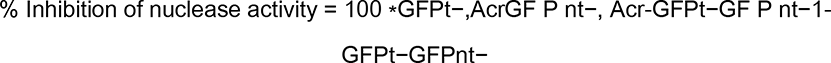

The inhibition of nuclease activity [%] is defined by the ratio of fluorescence between GFP targeting (GFPt-) and non-targeting (GFPnt-) Cas nucleases in the presence and absence of Acrs.

#### Quantification of SOS response

To measure the RecA-dependent SOS response, the nuclease-crRNA, the target/non-target plasmids (pCBS276/pCBS3578, kanamycin), and the reporter P*recA-gfp/no-gfp* (pCBS3611/pCBS3616, ampicillin) plasmids were transformed into *E. coli* BL21(AI) sequentially. Plasmids pCBS273 and pCBS3588 (chloramphenicol) were used to express SuCas12a2 and LbCas12a nucleases, respectively. When measuring the RecA-dependent SOS-response in the presence of LsCas13a, the nuclease expression plasmid pCBS361 (chloramphenicol) was used. The target/non-target plasmids pCBS2004/pCBS612 (ampicillin) and P*recA-gfp/no-gfp* plasmids pCBS2000/pCBS2002 (kanamycin) were used. First, the cells were grown LB medium with 0.2% glucose to inhibit expression of the nucleases and the crRNA. The bacteria were harvested from the overnight cultures (15 ml) by centrifugation at 5,000 g for 2 minutes and resuspended in fresh LB. Next, 200 µl of fresh LB medium were inoculated on 96 well plates with the resuspended bacteria from the overnight cultures. These cultures were grown in the presence of either chloramphenicol, kanamycin, and ampicillin, chloramphenicol and ampicillin, or no antibiotics. For induction of nuclease and crRNA expression 1 mM of IPTG and 0.2% L-arabinose were added.

The cultures were grown at 37°C with vigorous shaking. OD_600_ and fluorescence measurements (Excitation: 485/20, Emission: 528/20) were collected every 5 minutes on a BioTek Synergy H1 plate reader. Four biological replicates were measured per experimental condition. To determine if a change in fluorescence occurred as a result of nuclease targeting, first the background fluorescence collected for the cultures with the P*recA*-*no-gfp* plasmid (pCBS3616/pCBS2002) was subtracted from the values obtained for the cultures with the GFP-expressing plasmids for each time point (pCBS3611/pCBS2000). Next, the fluorescence values were divided by the OD_600_ values from the corresponding target and the non-target cultures. Statistical significance was determined using Welch’s t-test with unequal variance.

In parallel we performed a plasmid clearance assay with the washed overnight cultures (Extended Data Figure 15b), as described above. For the lowest plated dilution cultures at the OD_600_ of ∼0.1 were used.

#### Flow cytometry

For the flow cytometry measurements, *E. coli* BL21(AI) cells were sequentially electroporated with the nuclease-encoding and target/non-target plasmids. The SuCas12a2- and LbCas12a-expressing plasmids pCBS273 and pCBS3588 were used, respectively. Target plasmid pCBS273 and non-target plasmid pCBS3578 were used. For the experiments involving LsCas13a, nuclease-expression plasmid pCBS361 was used in combination with the target plasmid pCBS2004 and non-target plasmid pCBS612. Following plasmid transformation, the *E. coli* bacteria were recovered in SOC medium and grown overnight in LB with chloramphenicol, kanamycin, and 0.2% glucose. Next, the cells were harvested at 5,000 g for 2 minutes and resuspended in fresh LB. The resuspended bacteria were used to inoculate 15 ml cultures to the OD_600_ of ∼0.01. These cultures were grown at 37°C with 220 rpm shaking for 6 hours without antibiotics with 1 mM IPTG and 0.2% L-arabinose. Every 2 hours the OD_600_ of the cultures was measured and 500 µl samples were collected and centrifuged for 3 minutes at 5,000 g. The cell pellets were then resuspended in 1x PBS containing 2 µg/ml 4′,6-diamidino-2-phenylindole (DAPI, ThermoFisher Scientific, 62248). The resuspended cells were stained for 10 minutes in the dark, after which 10 µl were transferred into 240 µl of 1x PBS on a 96 well plate. DAPI fluorescence was measured with Cytoflow Novocyte Quanteon flow cytometer as emission in the Pacific Blue spectrum (455 nm). Data regarding the forward scatter (FSC) and the side scatter (SSC) were also collected.

The resulting data were analyzed in Python. First, clusters of bacteria which exhibit distinct FSC and Pacific Blue signals were identified using density-based spatial clustering of applications with noise (DBSCAN https://scikit-learn.org/stable/modules/generated/sklearn.cluster.DBSCAN.html). Next, the ratios of the Pacific Blue to the FSC signal for each data point and the percentage of the data points within each cluster were parsed from the clustering data. The resulting values were plotted in the form of balloon plots. 60,000 events per sample were analyzed.

#### Dead/Live staining

Dead and viable bacteria were estimated with the LIVE/DEAD BacLight™ Bacterial Viability and Counting Kit (Molecular Probes, L34856). The measurements were performed with Cytoflow Novocyte Quanteon flow cytometer. *E. coli* BL21(AI) bacteria were transformed with nuclease, crRNA, and either target or non-target expression plasmids. For expressing SuCas12a2 and LbCas12a with a target guide, plasmids pCBS273 and pCBS3588 were used, respectively. Target expression plasmid pCBS2004 and non-target expression plasmid pCBS612 were used. For expressing LsCas13a with a target and non-target guides, plasmids pCBS273 and pCBS3578 were used, respectively. Cultures containing combinations of nuclease-guide and target plasmids were grown for approximately 16 hours with 0.2% glucose inhibitor in four biological replicates. Afterwards, 1 ml of each culture was harvested at 5,000 g for 3 minutes. The resulting pellet was resuspended in 1 ml of fresh LB medium. 60 µl of this suspension were used to inoculate 20 ml of LB. There cultures were grown for 2 hours at 37°C with constant shaking at 220 rpm. The expression of the nucleases and the guides was induced with 0.2% arabinose and 0.01 mM IPTG. After 4 hours, OD600 of the cultures was measured. Volume of the cultures corresponding to the OD600 1.0 was collected and processed as described in the kit manual. Briefly, samples of the bacterial culture were spun in a centrifuge at 10,000 × g for 3 minutes to pellet the cells. The supernatant was removed and the pellet resuspended in 1 ml of 0.85% NaCl. As a control for the dead cells, spectate pellet was first resuspended in 300 μl 0.85% NaCl and then 700 μl 70% isopropyl alcohol (dead-cell suspension). The samples were incubated at room temperature for 60 minutes, with mixing every 15 minutes. Next, the samples were centrifuged at 10,000 × g for 3 minutes and washed in 1 ml 0.85% NaCl, followed by another centrifugation. Finally, the samples were resuspended min 0.5 ml of 0.85% NaCl. 1 ml of master mix for staining the cells contained 977 µl of 0.85% NaCl, 1.5 µl of Component A (3.34 mM SYTO 9 nucleic acid stain), 1.5 µl of Component B (30 mM propidium iodide(PI)), 10 µl of Component C (beads), and 10 µl of a sample. These reactions were incubated for 15 minutes at room temperature protected from light. Fluorescence was collected in the green (e.g., fluorescein for SYTO 9) and red (Texas Red for PI) channels. The dead cells in each sample were gated based on the dead-cell suspension control treated with isopropyl alcohol. The percentage of dead cells stained with PI was calculated from the total number of events without the beads. 50,000 events were counted per sample.

#### *In vivo* RNA degradation

Samples corresponding to 1 ml of culture at OD600 0.4 of grown for dead/live staining, as described above, were collected and centrifuged at 10,000 × g for 3 minutes. The resulting pellets were frozen in liquid nitrogen and stored −80°C until further processing. Total RNA was expected using 1.5 ml of trizol and 1.5 ml of ethanol with Direct-zol RNA Miniprep kit (R2051, Zymo), according to the manufacturer’s instructions. The RNA was further purified with the RNA Clean & Concentrator-5 kit (R1013, Zymo). 0.5 µg of RNA from each sample in 5 µl were combined with 2.5 µl of RNA loading dye, heated to 70°C for 10 minutes, and subsequently chilled on ice for 2 minutes. The used RNA High-Range ladder was also heat-treated. The denatured samples (5 µl) and the leader (3 µl) were loaded on a 1% TBE gel. The gel was run for 40 minutes at 120V. Afterwards, the gel was stained for 30 min in ethidium bromide, washed for 10 ml, and imaged.

#### Microscopy

For confocal microscopy, the cells were grown as for the flow cytometry above. In 2 hour intervals, 500 µl of each culture were collected and centrifuged at 5,000 g for 3 minutes. Next, the bacteria were diluted to approximately the same cell density and stained with 2 µg/ml of FM4-64 dye (ThermoFisher Scientific, T13320) and 1 µg/ml of DAPI (ThermoFisher Scientific, 62248). Imagining was done on a Leica DMi6000B TCS-SP5 II Inverted Confocal Microscope at 1,000x magnification.

## EXTENDED DATA FIGURES

**Extended Data Figure 1.**
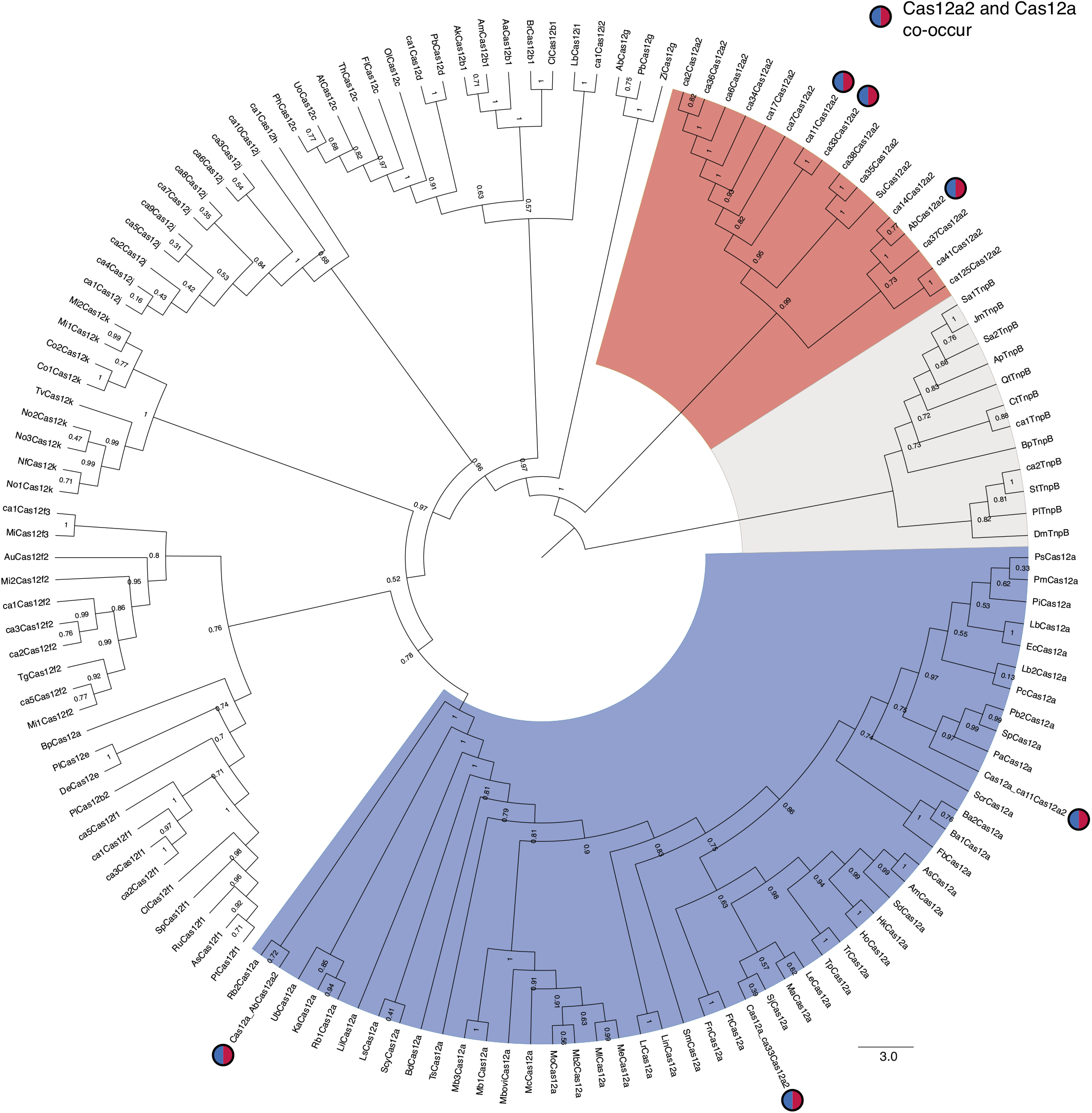
Maximum-likelihood phylogenetic inference of Cas12a2 and several other type V nucleases. The phylogenetic tree generated using amino acid sequences is an enlarged and annotated version of Figure 1a. The shaded regions designate Cas12a2 (red), Cas12a (blue), and TnpB (gray) orthologs. The blue and red filled circles indicate systems that contain both Cas12a2 and Cas12a.

**Extended Data Figure 2.**
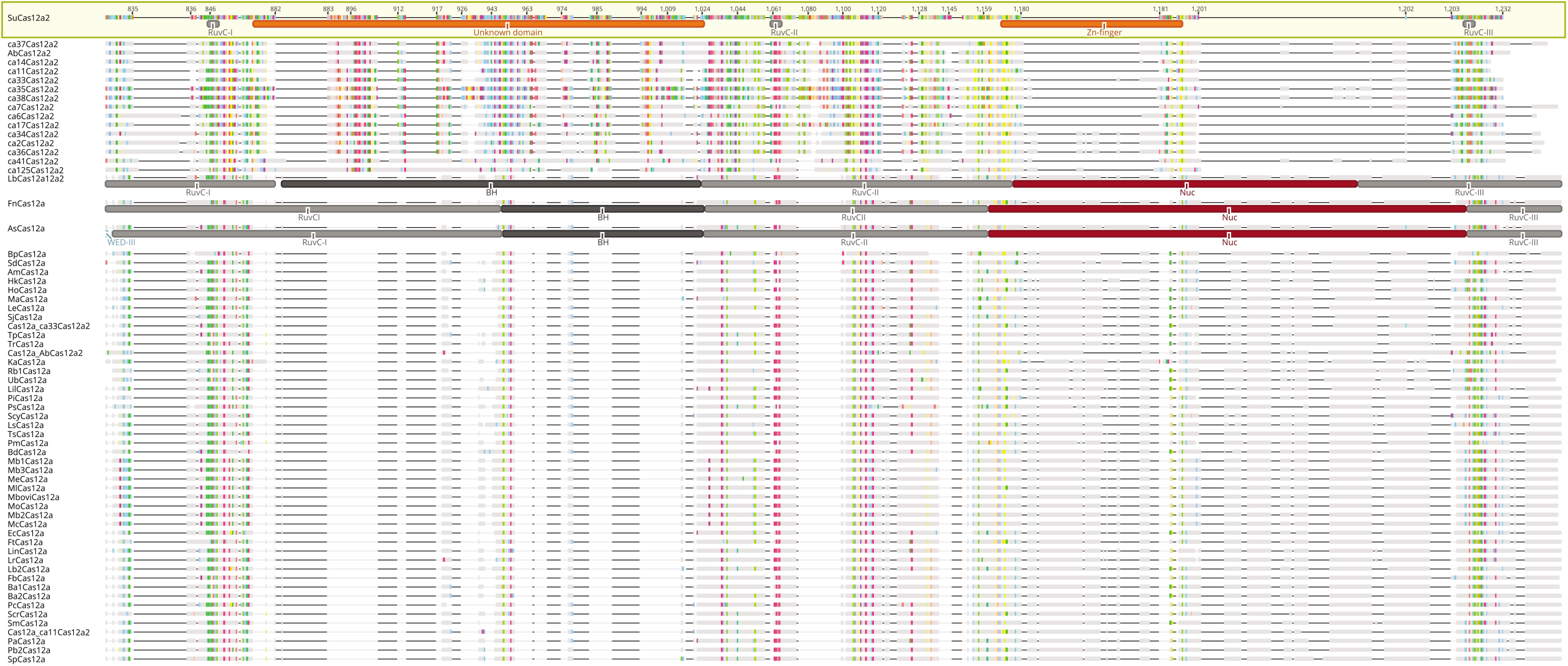
Fragment of an amino-acid sequence alignment containing the RuvC domain within Cas12a2 and Cas12a orthologs. The domains in FnCas12a^64^, LbCas12a^65^, and AsCas12a^66^ are based on crystal structures. The domains in SuCas12a2 were predicted using MOTIF search. The highlighted residues represent SuCas12a2 amino acids conserved in other Cas12a2 and Cas12a sequences. Numbering is shown in relation to SuCas12a2.

**Extended Data Figure 3.**
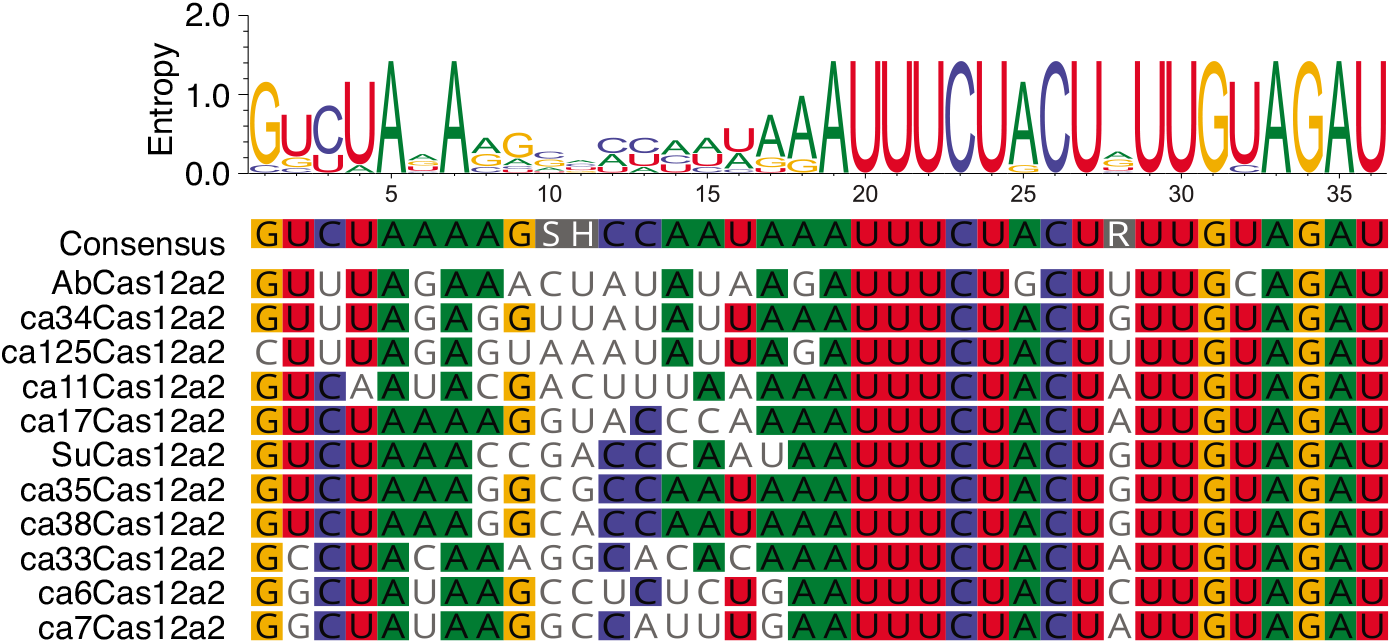
Nucleotide alignment of the direct repeats from CRISPR arrays associated with *cas12a2* genes. Flanking arrays could not be identified for some *cas12a2* genes.

**Extended Data Figure 4.**
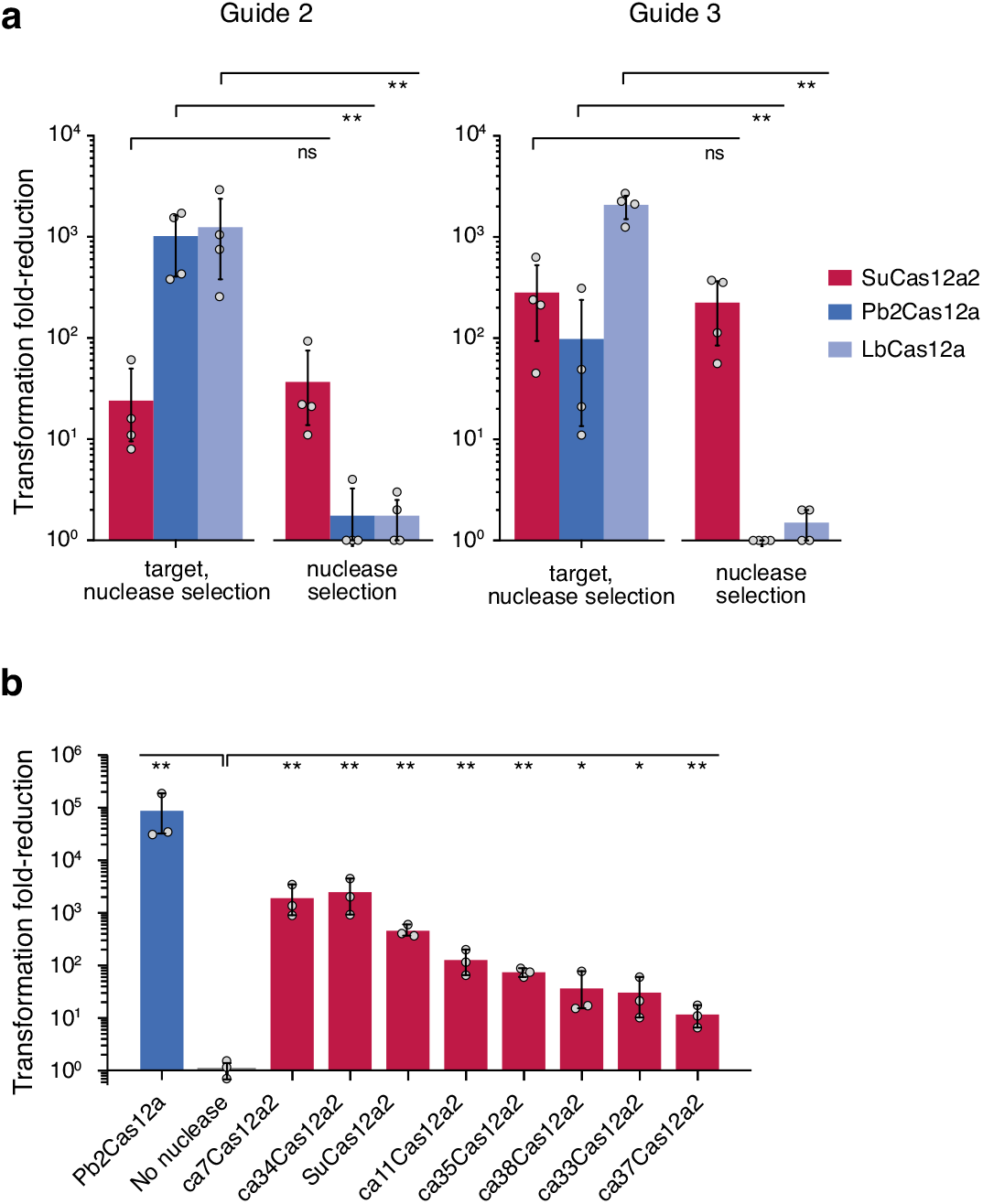
Transformation reduction by SuCas12a2 and other Cas12a2 orthologs, LbCas12a, and Pb2Cas12a in *E. coli* BL21(AI). (**a**) Different guide:target pairs tested under antibiotic selection of the nuclease and target plasmids or only the nuclease plasmid. (**b**) Different Cas12a2 orthologs tested under antibiotic selection of the nuclease and target plasmids. Error bars depict one standard deviation from the mean. Significance was calculated using Welch’s t-test (* - p < 0.05, ** - p < 0.005).

**Extended Data Figure 5.**
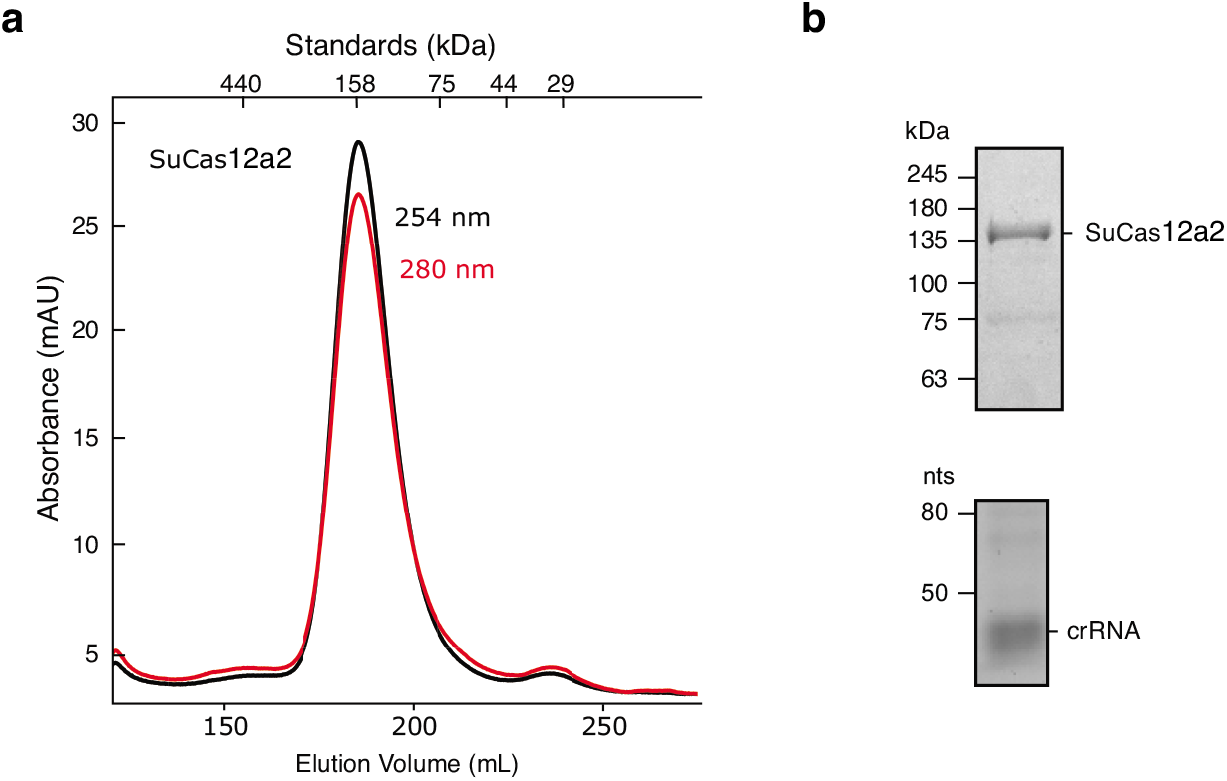
Purification of SuCas12a2. (**a**) Size exclusion chromatogram of purified SuCas12a2 bound to a co-expressed 3X crRNA guide over a Superdex 200 pg 26/600 column. Absorbance was measured at 254 nm and 280 nm. Molecular weight standards are indicated (top). (**b**) SDS-PAGE of purified SuCas12a2 + crRNA (top). Urea-PAGE gel of RNA acid-phenol-chloroform extracted from purified SuCas12a2 + crRNA (bottom).

**Extended Data Figure 6.**
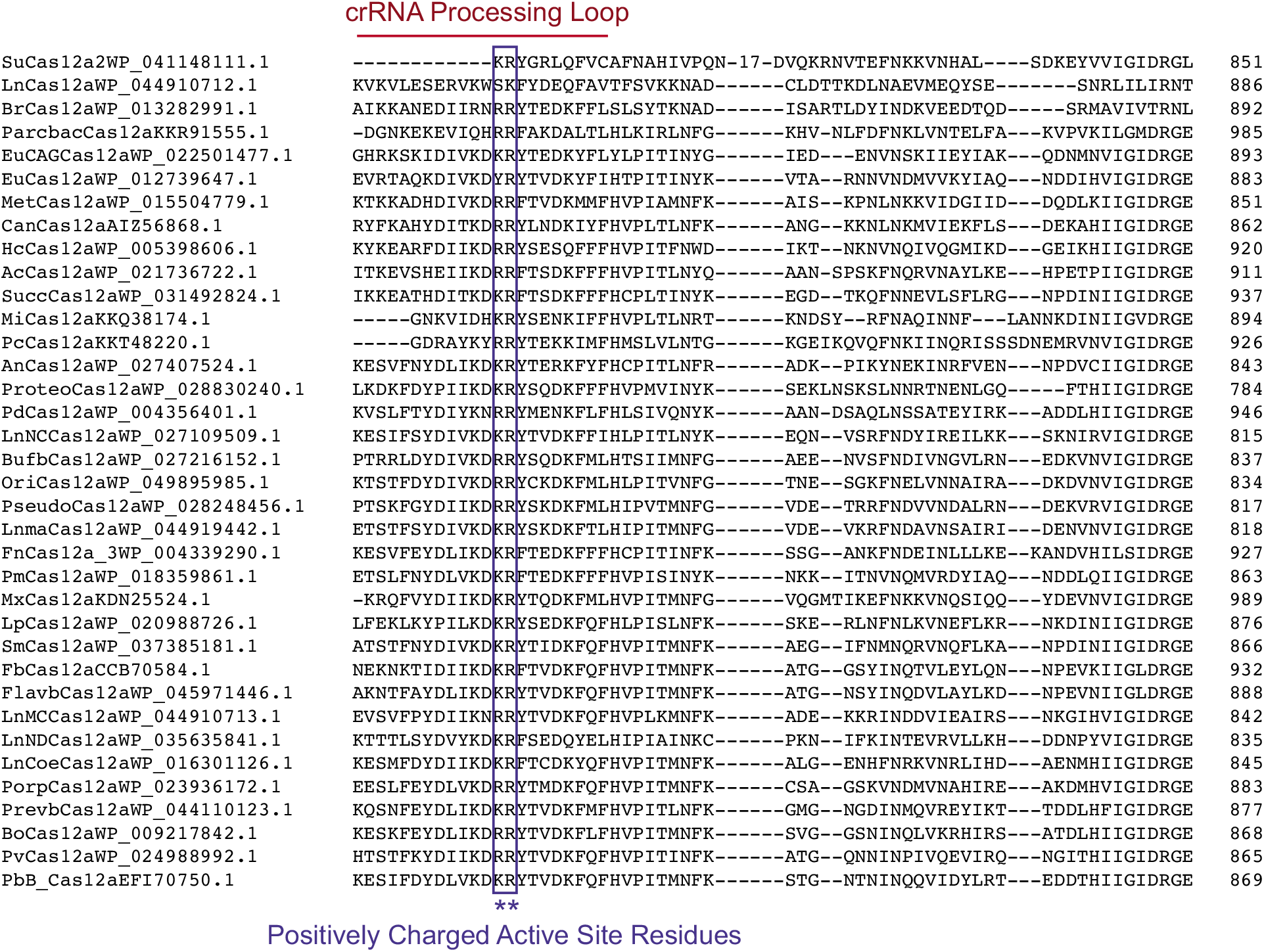
Clustal Omega amino acid sequence alignment of SuCas12a2 with Cas12a sequences. Cas12a amino acids located within the loop involved in crRNA-processing are indicated in red. Conserved positively charged residues involved in processing^21^ are indicated. Although the putative crRNA-processing loop of Cas12a2 is not as long the positively charged residues appear to be conserved.

**Extended Data Figure 7.**
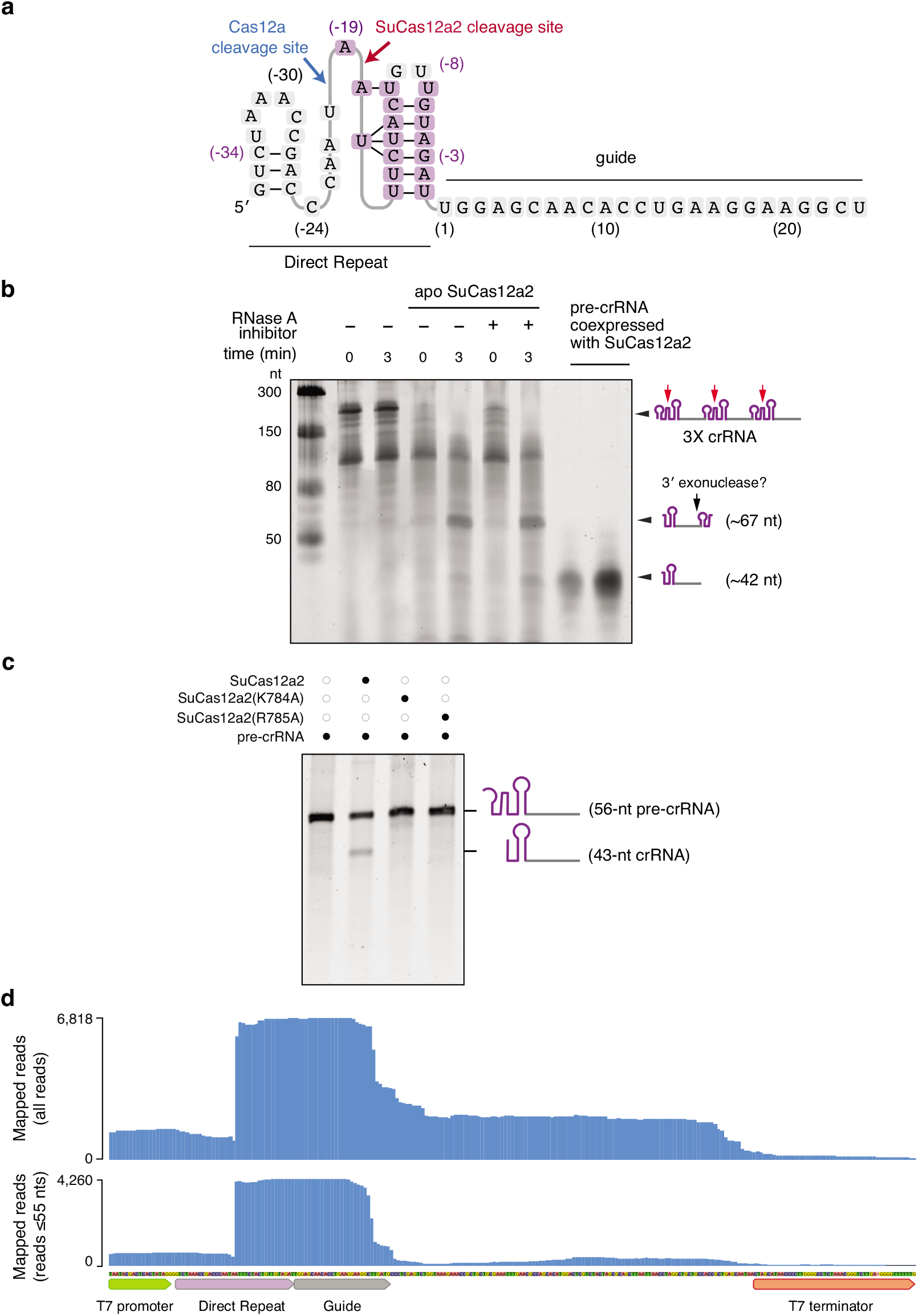
Pre-crRNA processing by SuCas12a2. (**a**) A diagram of the predicted secondary structure of the SuCas12a2 direct repeat. Cleavage sites of SuCas12a2 and Cas12a are indicated. (**b**) *In vitro* SuCas12a2-mediated cleavage of a pre-crRNA containing three direct repeats and three spacers (3X crRNA). Reactions containing apo-SuCas12a2 are indicated. Time points after mixing the cleavage reaction with apo-SuCas12a2 are indicated. The estimated sizes of the pre-crRNA and crRNAs after cleavage are indicated on the left. The last two lanes contain RNA extracted from SuCas12a2 co-expressed with a CRISPR array. The difference in size between the major crRNA band in the *in vitro* assay (∼ 67 nt) and the crRNA extracted from SuCas12a2 bound to *E. coli*-expressed crRNA (∼42) may be due to further trimming by 3′ exonucleases in the cell. (**c**) *In vitro* processing of a 56-nt pre-crRNA incubated for 60 minutes in the presence of apo-Cas12a2 or two mutants predicted to disrupt crRNA processing. (**d**) Sequencing coverage of the cDNA mapped to the crRNA locus in *E. coli* BL21(AI) expressing SuCas12a2. Coverage for all of the quality-filtered reads above 10 nts as well as reads between 10 and 55 nts mapped to the plus strand are shown.

**Extended Data Figure 8.**
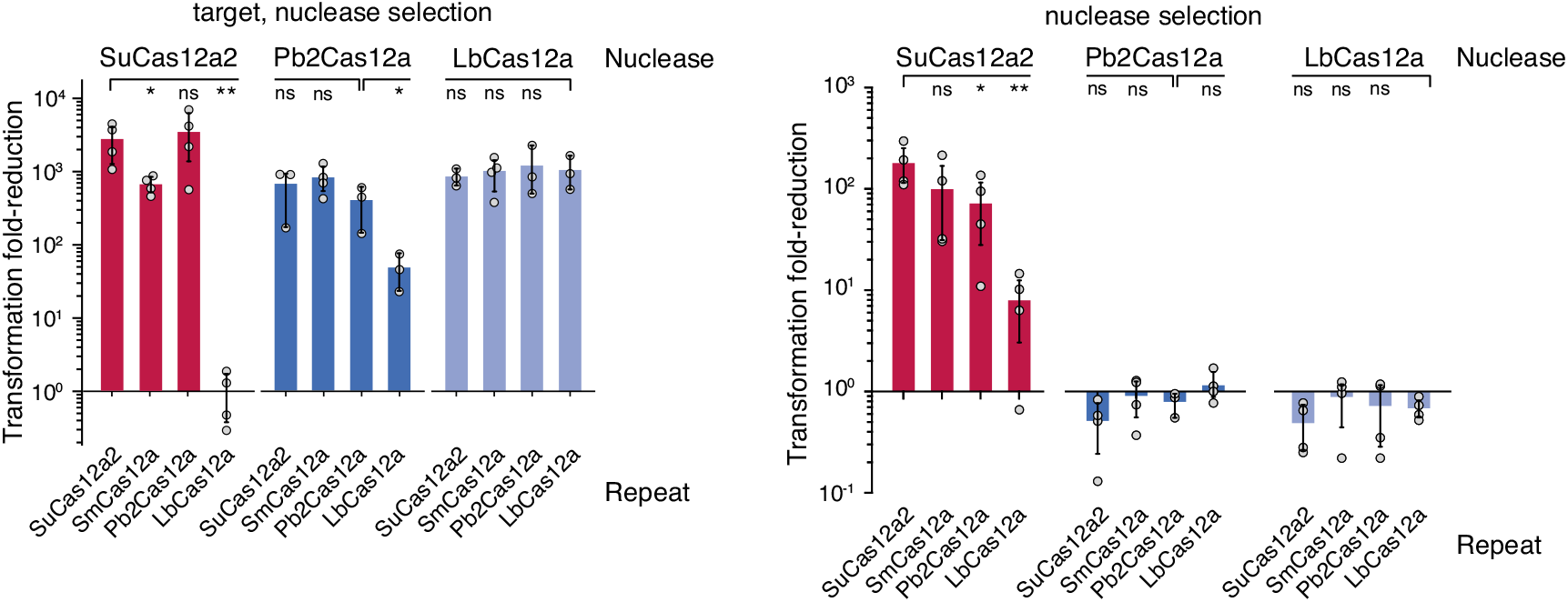
Cas12a and Cas12a2 can swap crRNA repeats. The indicated nuclease, repeat-encoding crRNA, and target were subjected to the traditional (left) and modified (right) plasmid interference assay in *E. coli*. Bar heights represent mean values from at least three independent experiments.

**Extended Data Figure 9.**
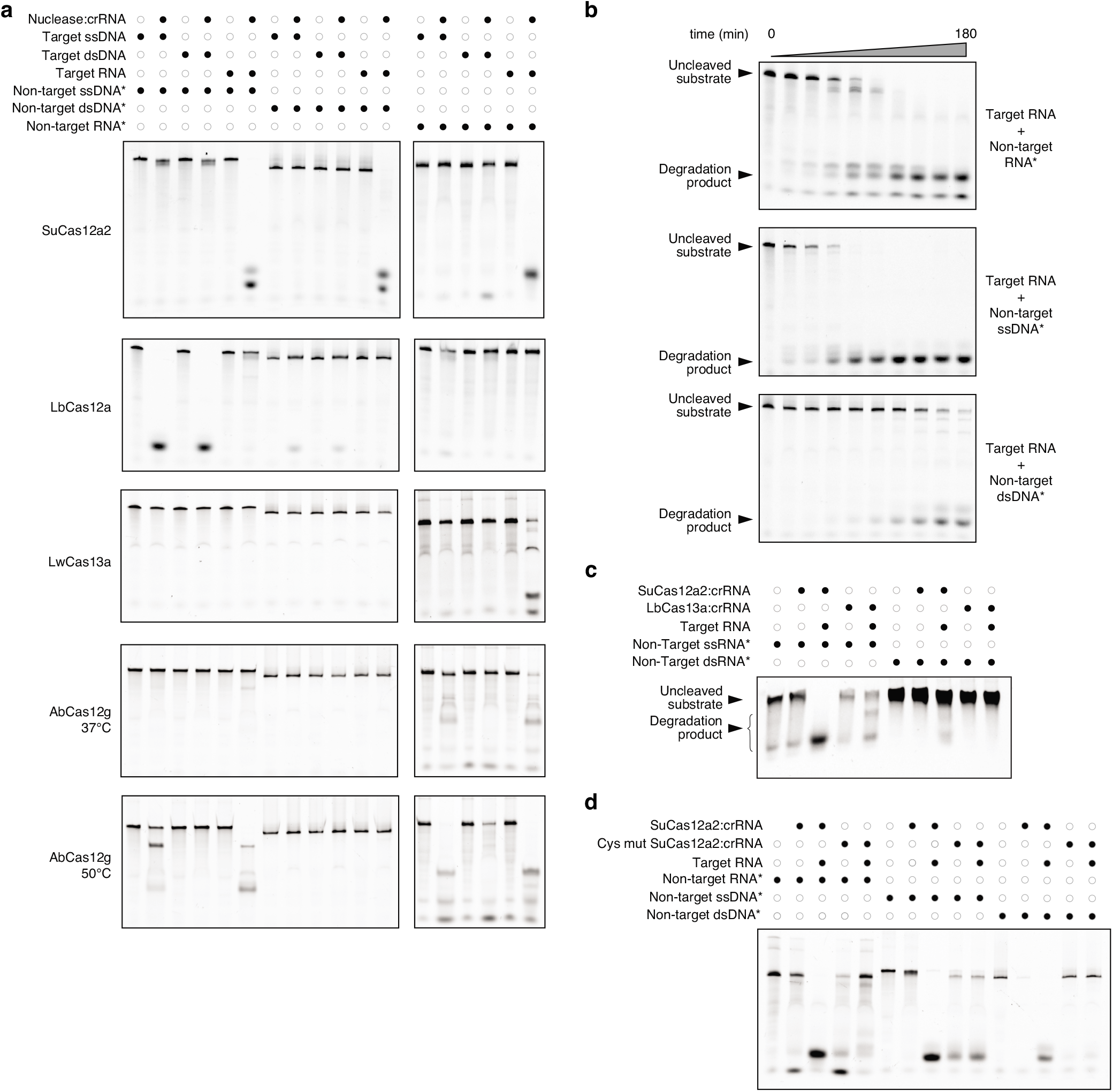
Properties of target recognition and collateral cleavage by SuCas12a2. (**a**) Non-specific collateral activities of SuCas12a2, LbCas12a, LwCas13a, and AbCas12g towards FAM-labeled non-target ssDNA, dsDNA, and RNA in the presence of target ssDNA, dsDNA, and RNA. All cleavage reactions were conducted at 37°C unless specified otherwise. AbCas12g was triggered by RNA and ssDNA, and it exhibited preferential collateral cleavage of RNA over ssDNA but no discernable cleavage of dsDNA. (**b**) SuCas12a2-mediated cleavage over time of FAM-labeled RNA (top), ssDNA (middle), and dsDNA (bottom) non-target substrates. These substrates are cleaved by SuCas12a2 through its non-specific collateral activity. (**c**) Electromobility shift assay indicating SuCas12a2 and LbCas13a do not indiscriminately degrade dsRNA. Uncleaved substrates and cleaved products are indicated. (**d**) Impact of mutating conserved cysteines within the predicted Zinc finger domain of SuCas12a2 on RNA-triggered collateral activity. The mutated cysteines within SuCas12a2 are C1170S, C1173S, C1188S and C1191S.

**Extended Data Figure 10.**
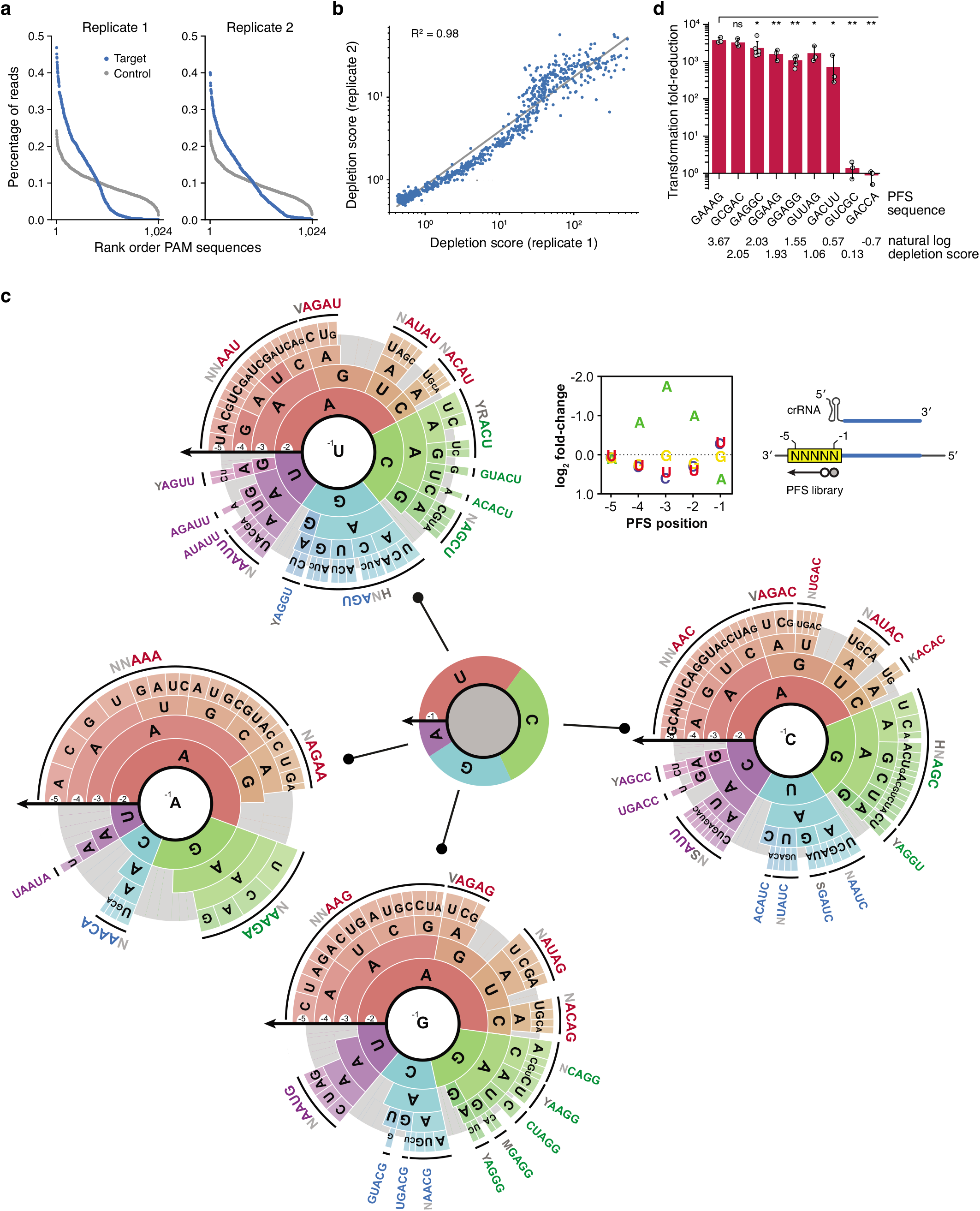
PFS depletion screen with SuCas12a2 in *E. coli* BL21(AI). (**a**) Sequencing coverage of the PFS libraries from the target and the control *E. coli* cultures. Data from two biological replicates are shown. (**b**) Correlation between the depletion scores obtained from the two replicate libraries. (**c**) The complex PFS profile recognized by SuCas12a2. See (ref. ^62^) for more information on interpreting PAM wheels. Given the complexity of the PFS profile, four different PAM wheels are shown based on each nucleotide at the −1 PFS position. (**d**) Validation of selected PFS sequences identified in the PFS screen with SuCas12a2. Bar heights represent mean values of at least three independent experiments. Error bars depict one standard deviation from the mean. Significance was calculated using Welch’s t-test (* - p < 0.05, ** - p < 0.005).

**Extended Data Figure 11.**
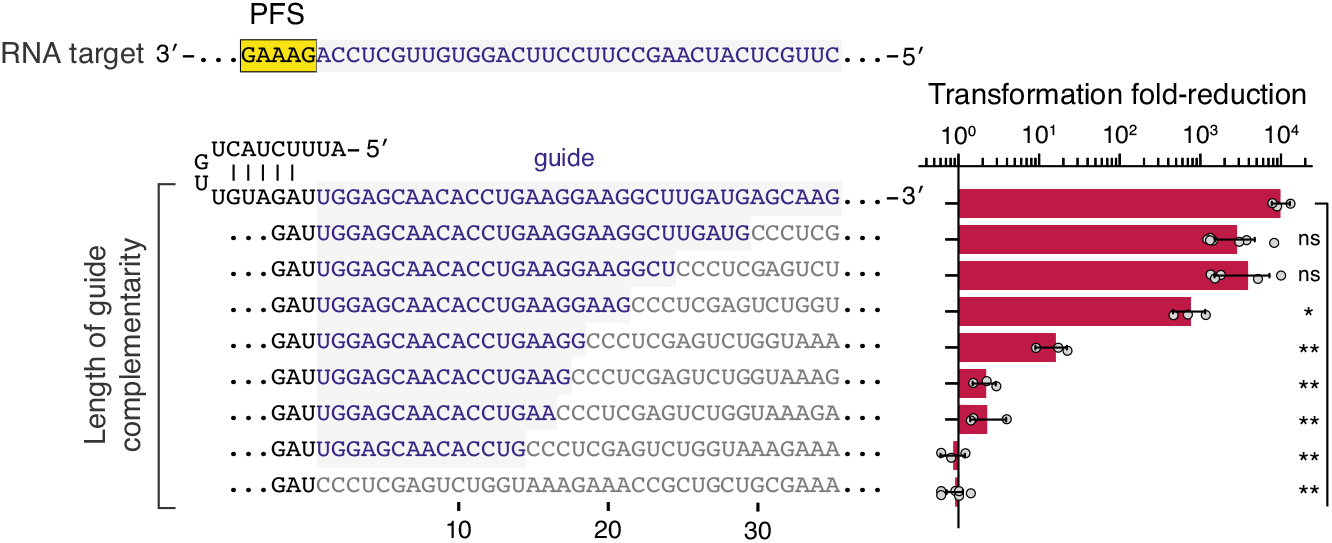
Impact of SuCas12a2 guide length on plasmid targeting in *E. coli* BL21(AI). The guide sequence is depicted with blue letters. The standard crRNA guide length based on crRNA processing (Extended Data Fig. 7) is 24 nts. Bar heights represent mean values from at least three independent experiments. Error bars depict one standard deviation from the mean. Significance was calculated using Welch’s t-test (* - p < 0.05, ** - p < 0.005).

**Extended Data Figure 12.**
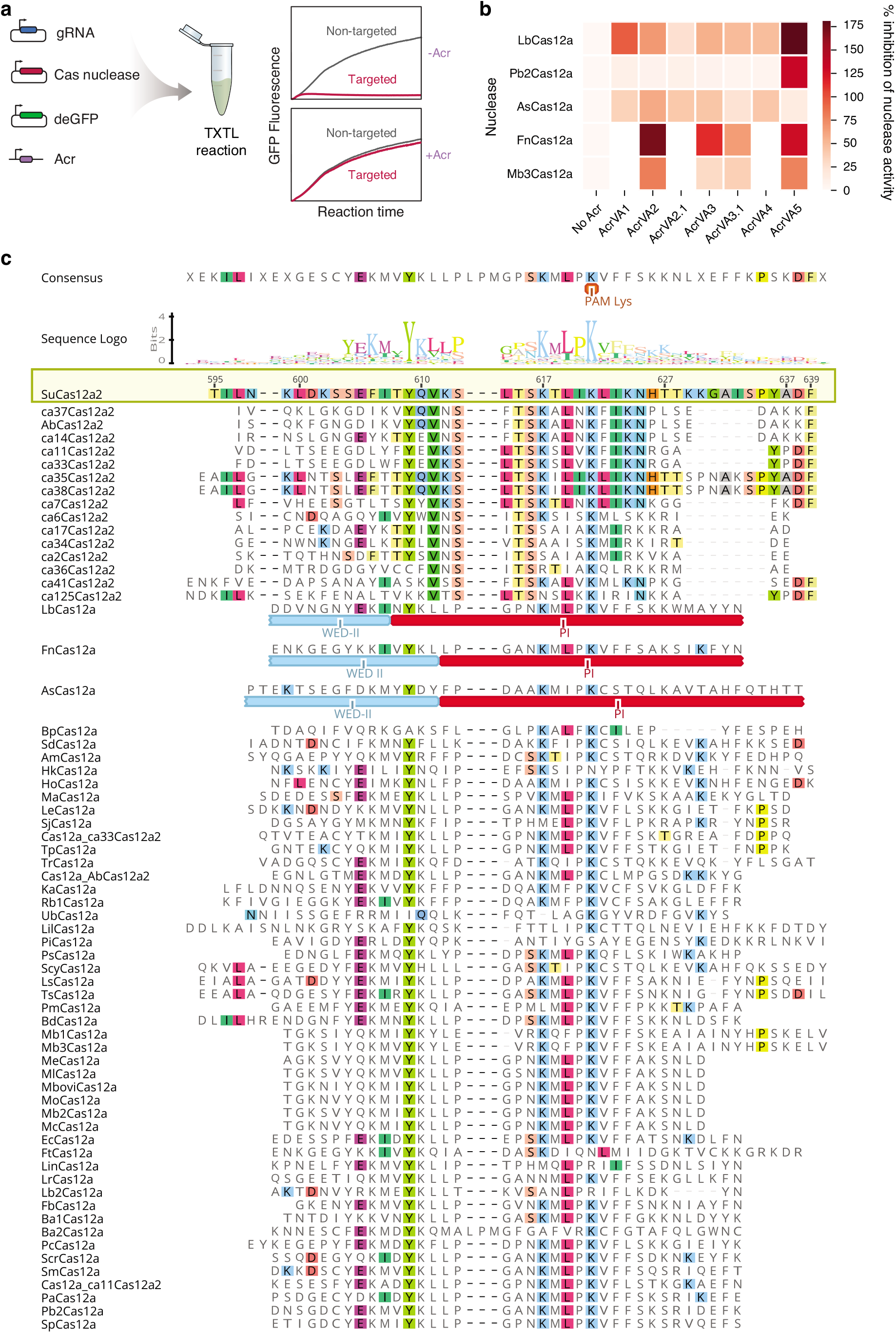
Verification of DNA targeting inhibition by AcrVA proteins in TXTL. (**a**) Schematic of the Acr targeting inhibition assay in TXTL. (**b**) Percent inhibition of Cas12a activity by the known type V-A anti-CRISPR proteins in TXTL. Inhibition above 100% reflects higher GFP levels for the target versus the non-target reaction in the presence of a given Acr. The mean of at least three biological replicates are shown. (**c**) Amino-acid sequence alignment of the locus containing the lysine residue acetylated by AcrVA5 in Cas12a^41^. The alignment shows that this residue is also present in Cas12a2 orthologs. The highlighted residues represent conserved amino acids relative to SuCas12a2. Numbering is shown in relation to SuCas12a2. The domains in FnCas12a^64^, LbCas12a^65^, and AsCas12a^66^ are based on crystal structures.

**Extended Data Figure 13.**
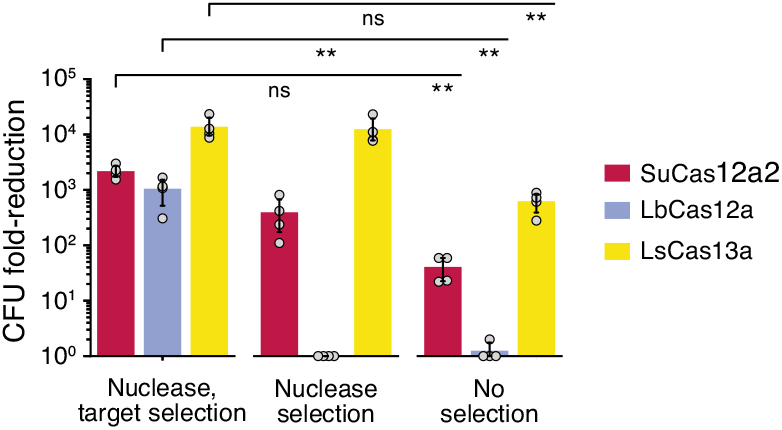
Reduction in colony-forming units (CFU) following nuclease and crRNA induction under different antibiotic selection conditions in *E. coli* BL21(AI). Bar heights represent mean values of at least three independent experiments. Error bars depict one standard deviation from the mean. Significance was calculated using Welch’s t-test (* - p < 0.05, ** - p < 0.005).

**Extended Data Figure 14.**
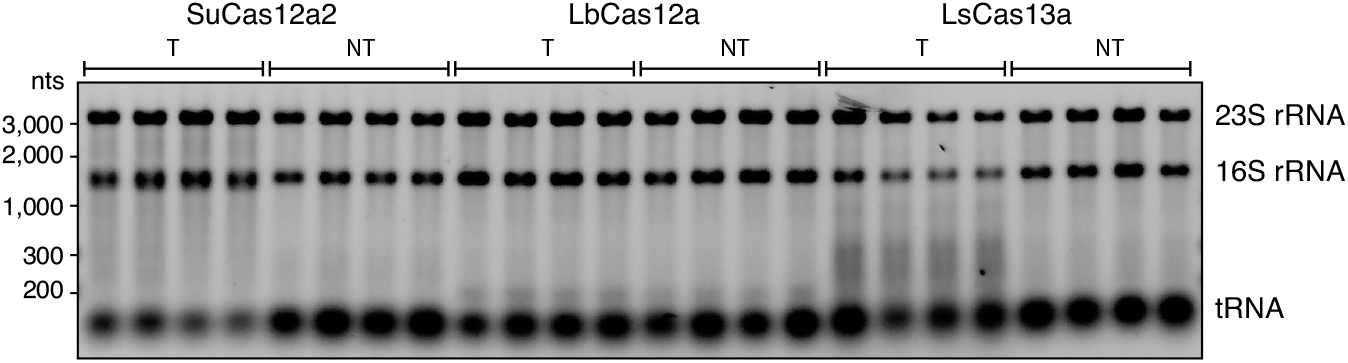
SuCas12a2 nuclease degrades cellular RNA. 1% agarose gel shows total RNA extracted with *E. coil* BL21(AI) expressing SuCas12a2, LbCas12a and LsCas13a under target (T) and non-target (NT) conditions. Expression of the nucleases and crRNA was induced with 10 nM IPTG and 0.2% arabinose for 2 hours. Individual wells for each condition represent biological replicates. Nucleotide sizes are based on a resolved RNA ladder.

**Extended Data Figure 15.**
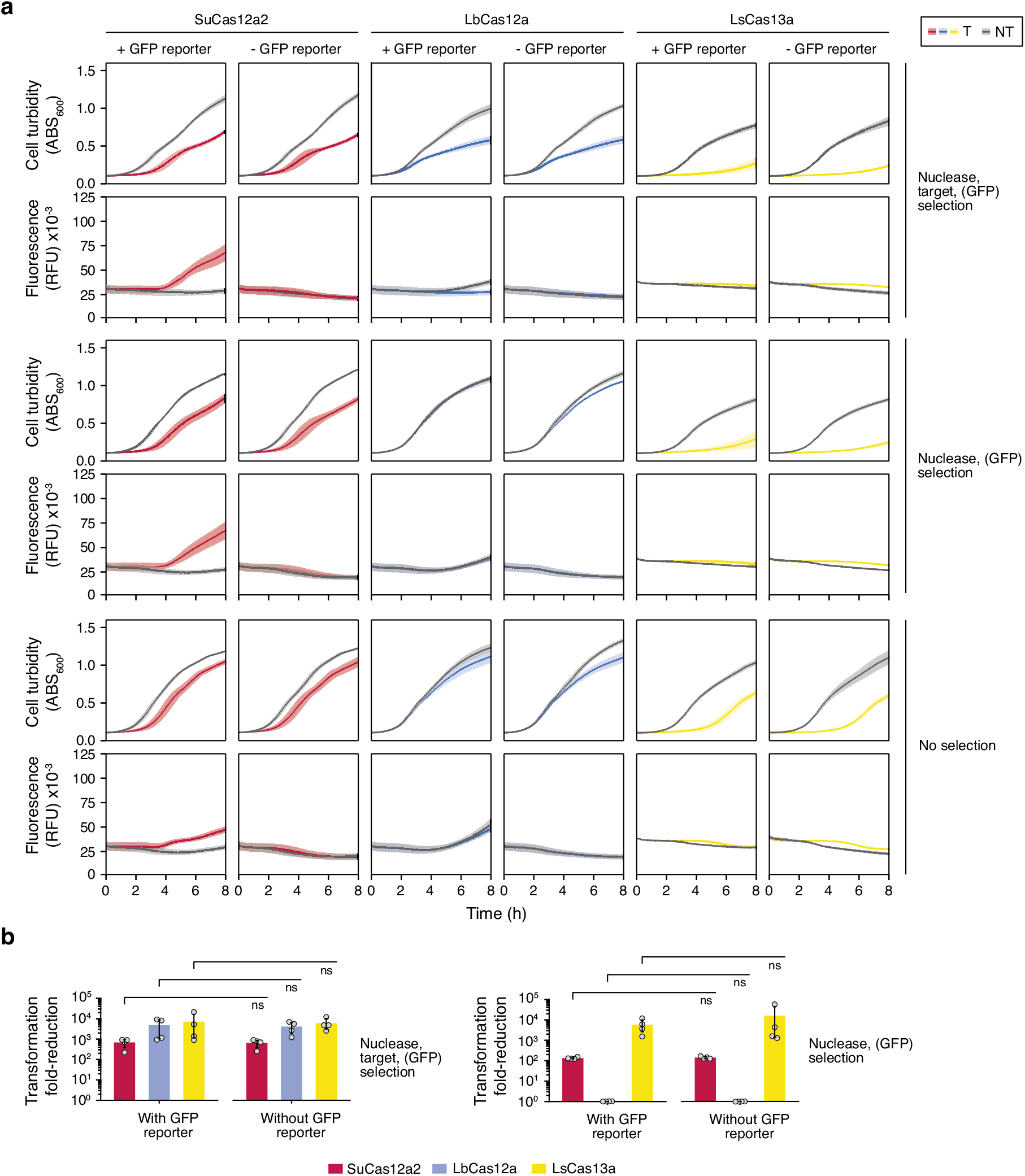
Inclusion of the SOS reporter construct does not perturb plasmid interference by the different Cas nucleases. (**a**) Turbidity and fluorescence time course measurements when assessing the SOS-responsive GFP reporter under different antibiotic selection conditions in *E. coli* BL21(AI). Darker bands represent the mean values of four independent experiments. The shaded areas depict one standard deviation from the mean. (**b**) Impact of the SOS-responsive GFP reporter on plasmid interference in *E. coli* BL21(AI) under different selection conditions. Bar heights represent mean values of four independent experiments. Error bars depict one standard deviation from the mean. Significance was calculated using Welch’s t-test (* - p < 0.05, ** - p < 0.005).

**Extended Data Figure 16.**
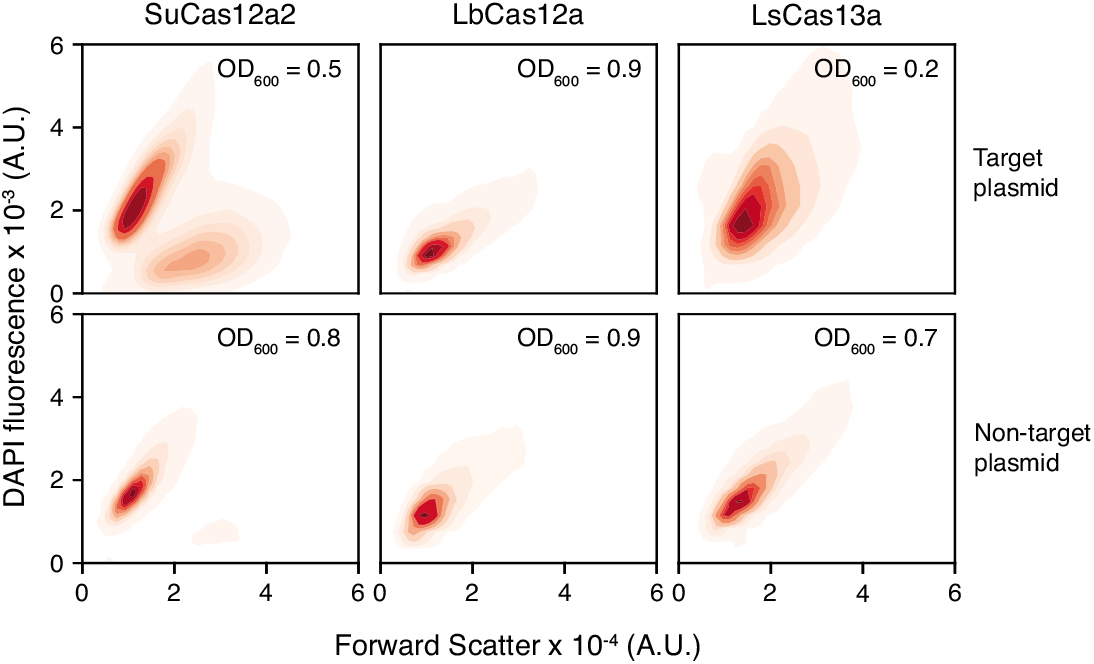
Flow cytometry analysis of the nuclease-expressing *E. coli* cells without antibiotic selection. The cultures were collected four hours after inoculation and induction of nuclease and crRNA. Prior to the analysis, the cells were stained with DAPI. The subpopulation with low DAPI and high forward scatter reflects elongated cells with reduced DNA content. Contour plots are representative of four independent experiments.

**Extended Data Figure 17.**
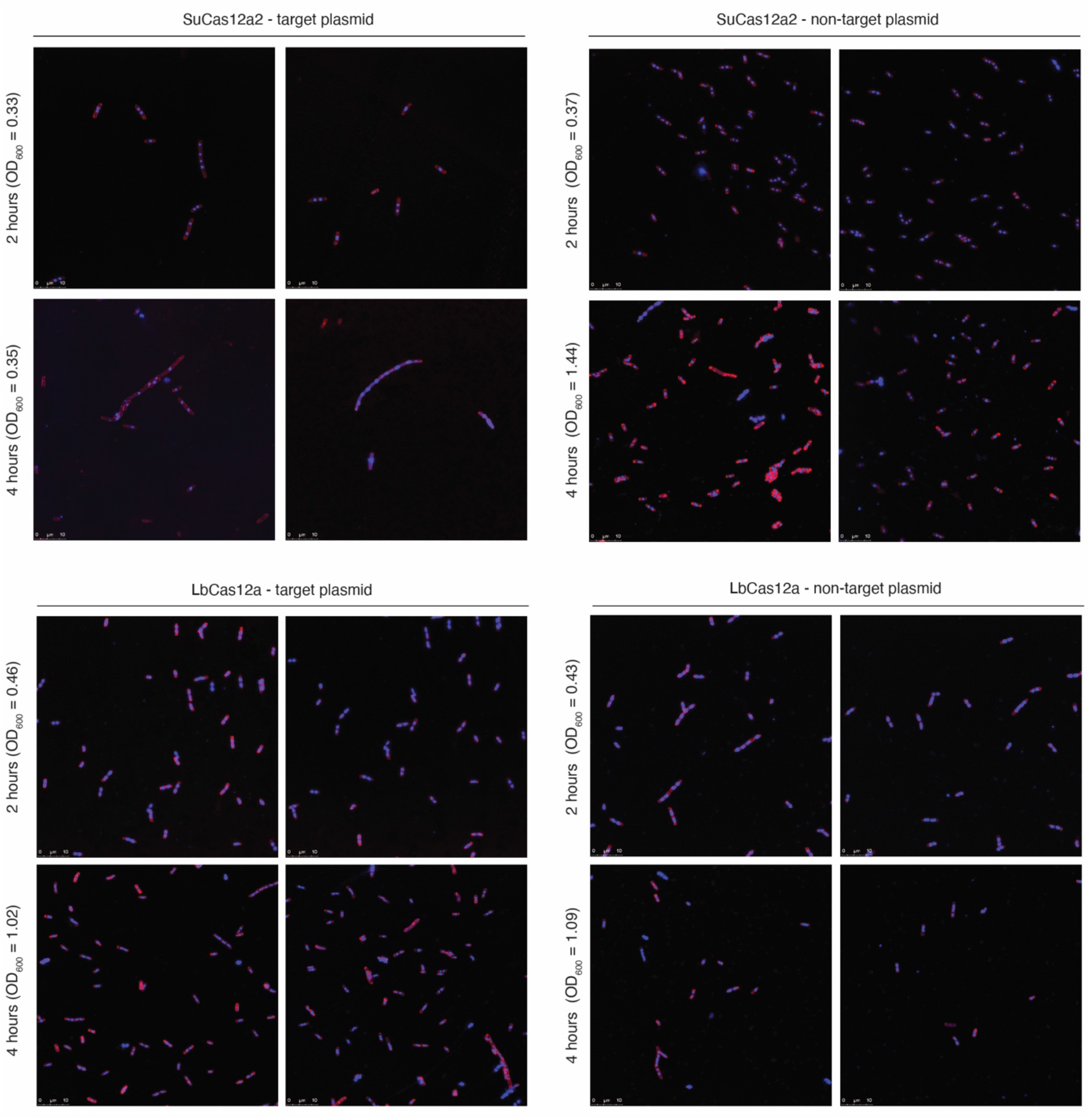
Confocal laser scanning micrographs of *E. coli* BL21(AI) transformed with SuCas12a2-crRNA and LbCas12-crRNA expression plasmids under target and non-target conditions in the absence of antibiotic selection. Extensive filamentation can be observed with the bacteria containing the target plasmid and expressing SuCas12a2. Within each overlaid image, fluorescent DNA (DAPI) staining is shown in blue and membrane (FM4-64) staining is shown in red. Paired images are different viewing fields from the same sample. Images are representative of two independent experiments.

**Extended Data Figure 18.**
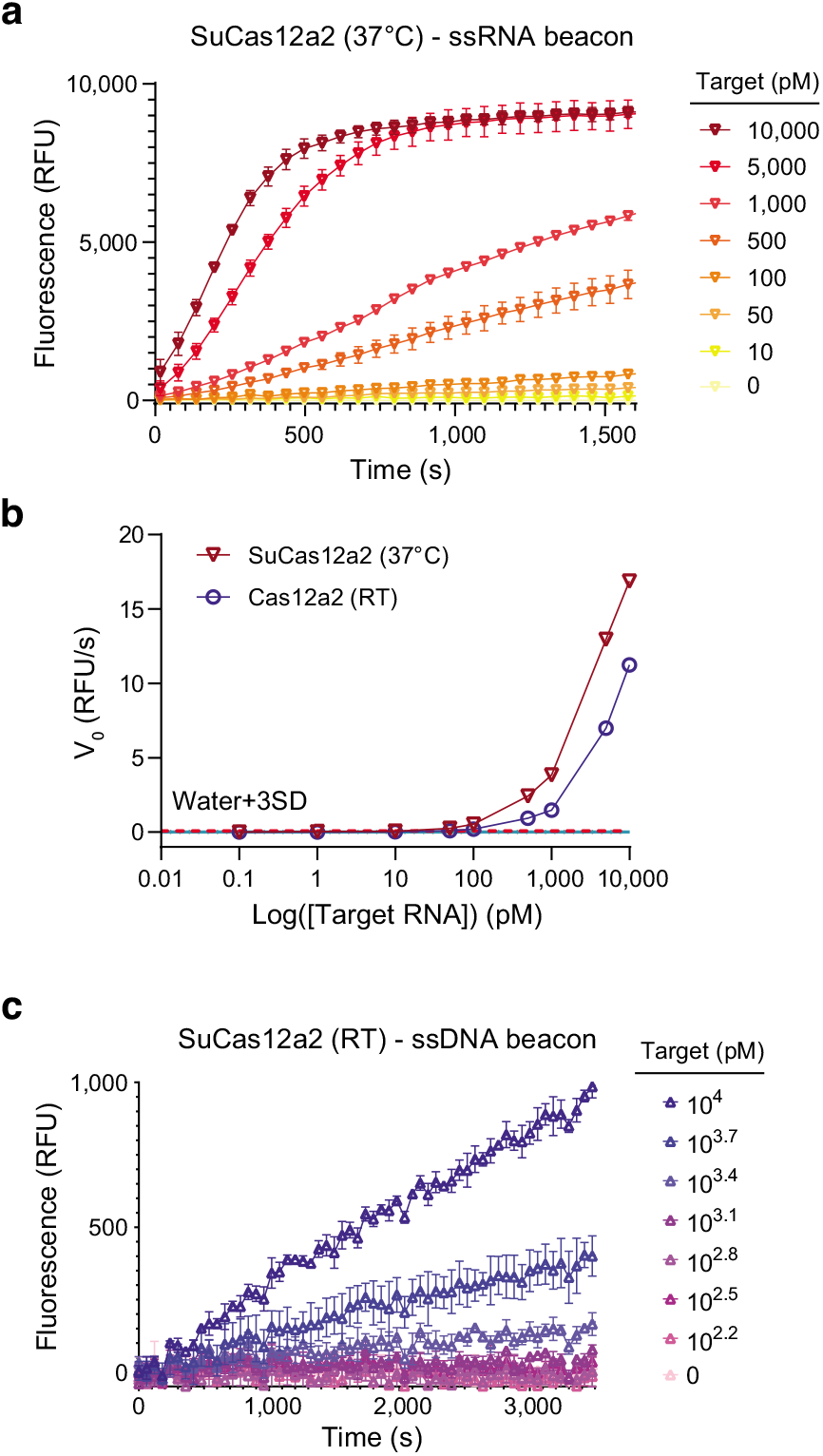
Determination of the limit of detection by Cas12a2 by velocity method^18^. (**a**) Progress curve for RNA activated cleavage of RNA beacon by Cas12a2 at 37°C. (b) Limit of detection of Cas12a2 using RNA beacon was determined using the velocity method. Velocities were obtained by regression analysis of the linear regions of the progress curves. The velocity method was used to determine all reported limits of detection. (**c**) Progress curve for RNA activated cleavage of ssDNA beacon by Cas12a2 at ambient temperature (RT).

## EXTENDED DATA TABLES

**Extended Data Table 1 |** Tabular data. See the Extended Data Table 1.xlsx file.

- Tab 1: List of strains used in this work.
- Tab 2: List of plasmids used in this work.
- Tab 3: List of key oligonucleotides, synthetic RNA, and dsDNA used in this work.

## EXTENDED DATA FILES

**Extended Data File 1 |** Fasta file listing Cas12a2 and other type V nucleases.

## REFERENCES

1. Lopatina, A., Tal, N. & Sorek, R. Abortive Infection: Bacterial Suicide as an Antiviral Immune Strategy. Annu. Rev. Virol. 7, 371–384 (2020).

2. Koonin, E. V. & Krupovic, M. Origin of programmed cell death from antiviral defense? Proc. Natl. Acad. Sci. 116, 16167–16169 (2019).

3. Meeske, A. J., Nakandakari-Higa, S. & Marraffini, L. A. Cas13-induced cellular dormancy prevents the rise of CRISPR-resistant bacteriophage. Nature 570, 241–245 (2019).

4. Rostøl, J. T. & Marraffini, L. Non-specific degradation of transcripts promotes plasmid clearance during type III-A CRISPR-Cas immunity. Nat. Microbiol. 4, 656–662 (2019).

5. Rostøl, J. T. et al. The Card1 nuclease provides defence during type III CRISPR immunity. Nature 590, 624–629 (2021).

6. VanderWal, A. R., Park, J.-U., Polevoda, B., Kellogg, E. H. & O’Connell, M. R. CRISPR-Csx28 forms a Cas13b-activated membrane pore required for robust CRISPR-Cas adaptive immunity. 2021.11.02.466367 (2021) doi:10.1101/2021.11.02.466367.

7. Kazlauskiene, M., Kostiuk, G., Venclovas, Č., Tamulaitis, G. & Siksnys, V. A cyclic oligonucleotide signaling pathway in type III CRISPR-Cas systems. Science 357, 605–609 (2017).

8. Niewoehner, O. et al. Type III CRISPR–Cas systems produce cyclic oligoadenylate second messengers. Nature 548, 543–548 (2017).

9. Makarova, K. S., Anantharaman, V., Grishin, N. V., Koonin, E. V. & Aravind, L. CARF and WYL domains: ligand-binding regulators of prokaryotic defense systems. Front. Genet. 5, (2014).

10. Gootenberg, J. S. et al. Nucleic acid detection with CRISPR-Cas13a/C2c2. Science 356, 438–442 (2017).

11. Abudayyeh, O. O. et al. C2c2 is a single-component programmable RNA-guided RNA-targeting CRISPR effector. Science 353, aaf5573 (2016).

12. Lau, R. K. et al. Structure and Mechanism of a Cyclic Trinucleotide-Activated Bacterial Endonuclease Mediating Bacteriophage Immunity. Mol. Cell 77, 723–733.e6 (2020).

13. Grüschow, S., Adamson, C. S. & White, M. F. Specificity and sensitivity of an RNA targeting type III CRISPR complex coupled with a NucC endonuclease effector. Nucleic Acids Res. 49, 13122–13134 (2021).

14. Chen, J. S. et al. CRISPR-Cas12a target binding unleashes indiscriminate single-stranded DNase activity. Science eaar6245 (2018) doi:10.1126/science.aar6245.

15. Marino, N. D., Pinilla-Redondo, R. & Bondy-Denomy, J. CRISPR-Cas12a targeting of ssDNA plays no detectable role in immunity. 2022.03.10.483831 (2022) doi:10.1101/2022.03.10.483831.

16. Zetsche, B. et al. Cpf1 Is a Single RNA-Guided Endonuclease of a Class 2 CRISPR-Cas System. Cell 163, 759–771 (2015).

17. Jinek, M. et al. A Programmable Dual-RNA–Guided DNA Endonuclease in Adaptive Bacterial Immunity. Science 337, 816–821 (2012).

18. Huyke, D. A. et al. Fundamental limits of amplification-free CRISPR-Cas12 and Cas13 diagnostics. http://biorxiv.org/lookup/doi/10.1101/2022.01.31.478567 (2022) doi:10.1101/2022.01.31.478567.

19. Makarova, K. S., Wolf, Y. I. & Koonin, E. V. Classification and Nomenclature of CRISPR-Cas Systems: Where from Here? CRISPR J. 1, 325–336 (2018).

20. Bravo, J. P. K. et al. Large-scale structural rearrangements unleash indiscriminate nuclease activity by CRISPR-Cas12a2. *Be Submitted*.

21. Fonfara, I., Richter, H., Bratovič, M., Rhun, A. L. & Charpentier, E. The CRISPR-associated DNA-cleaving enzyme Cpf1 also processes precursor CRISPR RNA. Nature 532, 517–521 (2016).

22. Swarts, D. C., van der Oost, J. & Jinek, M. Structural Basis for Guide RNA Processing and Seed-Dependent DNA Targeting by CRISPR-Cas12a. Mol. Cell 66, 221–233.e4 (2017).

23. East-Seletsky, A. et al. Two Distinct RNase Activities of CRISPR-C2c2 Enable Guide RNA Processing and RNA Detection. Nature 538, 270–273 (2016).

24. Samai, P. et al. Co-transcriptional DNA and RNA Cleavage during Type III CRISPR-Cas Immunity. Cell 161, 1164–1174 (2015).

25. Yan, W. X. et al. Functionally diverse type V CRISPR-Cas systems. Science 363, 88–91 (2019).

26. Marraffini, L. A. & Sontheimer, E. J. Self vs. non-self discrimination during CRISPR RNA-directed immunity. Nature 463, 568–571 (2010).

27. Estrella, M. A., Kuo, F.-T. & Bailey, S. RNA-activated DNA cleavage by the Type III-B CRISPR–Cas effector complex. Genes Dev. 30, 460–470 (2016).

28. Murugan, K., Seetharam, A. S., Severin, A. J. & Sashital, D. G. CRISPR-Cas12a has widespread off-target and dsDNA-nicking effects. J. Biol. Chem. 295, 5538–5553 (2020).

29. Marshall, R. et al. Rapid and Scalable Characterization of CRISPR Technologies Using an E. coli Cell-Free Transcription-Translation System. Mol. Cell 69, 146–157.e3 (2018).

30. Elmore, J. R. et al. Bipartite recognition of target RNAs activates DNA cleavage by the Type III-B CRISPR–Cas system. Genes Dev. 30, 447–459 (2016).

31. Kazlauskiene, M., Tamulaitis, G., Kostiuk, G., Venclovas, Č. & Siksnys, V. Spatiotemporal Control of Type III-A CRISPR-Cas Immunity: Coupling DNA Degradation with the Target RNA Recognition. Mol. Cell 62, 295–306 (2016).

32. Liu, T. Y., Iavarone, A. T. & Doudna, J. A. RNA and DNA Targeting by a Reconstituted Thermus thermophilus Type III-A CRISPR-Cas System. PLOS ONE 12, e0170552 (2017).

33. Han, W. et al. A type III-B CRISPR-Cas effector complex mediating massive target DNA destruction. Nucleic Acids Res. 45, 1983–1993 (2017).

34. Wang, B. et al. Structural basis for self-cleavage prevention by tag:anti-tag pairing complementarity in type VI Cas13 CRISPR systems. Mol. Cell 81, 1100–1115.e5 (2021).

35. Wiedenheft, B. et al. RNA-guided complex from a bacterial immune system enhances target recognition through seed sequence interactions. Proc. Natl. Acad. Sci. 108, 10092–10097 (2011).

36. Semenova, E. et al. Interference by clustered regularly interspaced short palindromic repeat (CRISPR) RNA is governed by a seed sequence. Proc. Natl. Acad. Sci. U. S. A. 108, 10098–10103 (2011).

37. Liu, L. et al. Two Distant Catalytic Sites Are Responsible for C2c2 RNase Activities. Cell 168, 121–134.e12 (2017).

38. Watters, K. E., Fellmann, C., Bai, H. B., Ren, S. M. & Doudna, J. A. Systematic discovery of natural CRISPR-Cas12a inhibitors. Science 362, 236–239 (2018).

39. Knott, G. J. et al. Broad-spectrum enzymatic inhibition of CRISPR-Cas12a. Nat. Struct. Mol. Biol. 26, 315–321 (2019).

40. Marino, N. D. et al. Discovery of widespread type I and type V CRISPR-Cas inhibitors. Science 362, 240–242 (2018).

41. Dong, L. et al. An anti-CRISPR protein disables type V Cas12a by acetylation. Nat. Struct. Mol. Biol. 26, 308–314 (2019).

42. Little, J. W. & Mount, D. W. The SOS regulatory system of Escherichia coli. Cell 29, 11–22 (1982).

43. Janion, C. Some aspects of the SOS response system--a critical survey. Acta Biochim. Pol. 48, 599–610 (2001).

44. Chen, Z., Lu, M., Zou, D. & Wang, H. An E. coli SOS-EGFP biosensor for fast and sensitive detection of DNA damaging agents. J. Environ. Sci. 24, 541–549 (2012).

45. Vercoe, R. B. et al. Cytotoxic Chromosomal Targeting by CRISPR/Cas Systems Can Reshape Bacterial Genomes and Expel or Remodel Pathogenicity Islands. PLOS Genet. 9, e1003454 (2013).

46. Cui, L. & Bikard, D. Consequences of Cas9 cleavage in the chromosome of Escherichia coli. Nucleic Acids Res. 44, 4243–4251 (2016).

47. Caliando, B. J. & Voigt, C. A. Targeted DNA degradation using a CRISPR device stably carried in the host genome. Nat. Commun. 6, 6989 (2015).

48. Nemudraia, A. et al. Sequence-specific capture and concentration of viral RNA by type III CRISPR system enhances diagnostic. Res. Sq. rs.3.rs-1466718 (2022) doi:10.21203/rs.3.rs-1466718/v1.

49. Silas, S. et al. Type III CRISPR-Cas systems can provide redundancy to counteract viral escape from type I systems. eLife 6, e27601 (2017).

50. Liu, L. et al. The Molecular Architecture for RNA-Guided RNA Cleavage by Cas13a. Cell 170, 714–726.e10 (2017).

51. Sievers, F. et al. Fast, scalable generation of high-quality protein multiple sequence alignments using Clustal Omega. Mol. Syst. Biol. 7, 539 (2011).

52. Kozlov, A. M., Darriba, D., Flouri, T., Morel, B. & Stamatakis, A. RAxML-NG: a fast, scalable and user-friendly tool for maximum likelihood phylogenetic inference. Bioinform. (2019) doi:10.1093/bioinformatics/btz305.

53. Kelley, L. A., Mezulis, S., Yates, C. M., Wass, M. N. & Sternberg, M. J. E. The Phyre2 web portal for protein modeling, prediction and analysis. Nat. Protoc. 10, 845–858 (2015).

54. Söding, J., Biegert, A. & Lupas, A. N. The HHpred interactive server for protein homology detection and structure prediction. Nucleic Acids Res. 33, W244–248 (2005).

55. Guzman, L. M., Belin, D., Carson, M. J. & Beckwith, J. Tight regulation, modulation, and high-level expression by vectors containing the arabinose PBAD promoter. J. Bacteriol. 177, 4121–4130 (1995).

56. Studier, F. W. Protein production by auto-induction in high density shaking cultures. Protein Expr. Purif. 41, 207–234 (2005).

57. Lapinaite, A., et al. DNA capture by a CRISPR-Cas9–guided adenine base editor. Science 369, 566–571 (2020).

58. Harris, C. J., Molnar, A., Müller, S. Y. & Baulcombe, D. C. FDF-PAGE: a powerful technique revealing previously undetected small RNAs sequestered by complementary transcripts. Nucleic Acids Res. 43, 7590–7599 (2015).

59. Li, Z., Zhang, H., Xiao, R., Han, R. & Chang, L. Cryo-EM structure of the RNA-guided ribonuclease Cas12g. Nat. Chem. Biol. 17, 387–393 (2021).

60. Bushnell, B., Rood, J. & Singer, E. BBMerge – Accurate paired shotgun read merging via overlap. PLOS ONE 12, e0185056 (2017).

61. Green, M. R., Sambrook, J. & Sambrook, J. Molecular cloning: a laboratory manual. (Cold Spring Harbor Laboratory Press, 2012).

62. Leenay, R. T. et al. Identifying and Visualizing Functional PAM Diversity across CRISPR-Cas Systems. Mol. Cell 62, 137–147 (2016).

63. Wandera, K. G. et al. An enhanced assay to characterize anti-CRISPR proteins using a cell-free transcription-translation system. Methods San Diego Calif 172, 42–50 (2020).

64. Swarts, D. C. & Jinek, M. Mechanistic Insights into the cis-and trans-Acting DNase Activities of Cas12a. Mol. Cell 73, 589–600.e4 (2019).

65. Dong, D. et al. The crystal structure of Cpf1 in complex with CRISPR RNA. Nature 532, 522–526 (2016).

66. Yamano, T. et al. Crystal Structure of Cpf1 in Complex with Guide RNA and Target DNA. Cell 165, 949–962 (2016).

